# A *TP53* Intron-Derived Non-Canonical Protein Promotes Tumor Survival

**DOI:** 10.1101/2025.09.04.674312

**Authors:** Louis A. Cadena, Kevin Lee Luo, Sofia M. Fernandez, Cristian A. Otero-Perez, Yu-Wen Huang, Ruiying Zhao, Dung-Fang Lee, Angela H. Ting, Xin Hu, Agata Sekrecka, Michal Sekrecki, Sydney X.-L. Lu, Sanghyun Kim, Fengxi Ye, Mingjian J. You, Shih-Han Lee

**Affiliations:** Department of Genetics, The University of Texas MD Anderson Cancer Center, Houston, TX 77030, USA; Department of Integrative Biology and Pharmacology, McGovern Medical School, The University of Texas Health Science Center at Houston, Houston, TX 77030, USA; Department of Epigenetics and Molecular Carcinogenesis, The University of Texas MD Anderson Cancer Center, Houston, TX 77030, USA; Department of Thoracic/Head and Neck Medical Oncology, The University of Texas MD Anderson Cancer Center, Houston, TX, 77030, USA; Division of Hematology, Department of Medicine, Stanford University School of Medicine, Stanford, CA, USA; Department of Hematopathology, Division of Pathology and Laboratory Medicine, The University of Texas MD Anderson Cancer Center, Houston, TX 77030, USA; The University of Texas MD Anderson Cancer Center, UTHealth Houston Graduate School of Biomedical Sciences, Houston, TX 77030, USA; Department of Pathology, City of Hope National Medical Center, Duarte, CA, USA

**Author notes:** ^9^These authors contributed equally to this work (Louis A. Cadena, Kevin Lee Luo).

**Keywords:** cleavage and polyadenylation, intronic polyadenylation, RNA isoforms, non-canonical protein, leukemia

## Abstract

During transcription, alternative cleavage and polyadenylation (APA) of transcripts produces mRNAs that differ in their 3’ ends. We previously found that leukemia cells cleave and polyadenylate transcripts within introns (intronic A<u>PA</u>, IPA) to generate truncated RNA isoforms, which perturb the expression and function of corresponding genes. In addition to protein-coding RNA, we found that leukemia cells express numerous putative long non-coding RNAs generated by IPA, with most uncharacterized. Here, we identify an IPA isoform encoded within intron 1 of the *TP53* tumor suppressor gene that is translated into a non-canonical protein termed IPADP1 (<u>IPA</u>-<u>d</u>erived <u>p</u>rotein 1). IPADP1 expression was detected in patient leukemia cells and human blood cancer cell lines. We uncovered an oncogenic role for IPADP1 in enhancing cancer cell survival during drug treatments, as well as promoting tumor formation *in vivo*. Using proteomic and functional studies, our data suggest that IPADP1 increases cell viability by regulating apoptotic signaling, cell cycle, and DNA damage responses. Furthermore, we observed that the expression of *p53-IPA* is significantly associated with *SF3B1* mutations in leukemia patients, suggesting mechanisms for the genesis of this isoform. Our study demonstrates that an IPA isoform could have a function different from its host gene.

## INTRODUCTION

The role of RNA processing has been well-recognized in tumorigenesis and paves avenues for the discovery of mechanisms critical for malignant transformation^1,2^. During transcription, alternative cleavage and polyadenylation (APA) is used in a context and cell-type dependent manner to produce transcripts that differ in their 3’ ends without changing DNA sequences^3–6^. APA events are pervasive in normal and malignant cells, with approximately 70% of human genes being regulated by APA^1,3,4^.

It is known that APA can occur at multiple sites throughout a gene^1,3–5^. When cleavage and polyadenylation signals are located within an intron (intronic APA, IPA), truncated RNA isoforms are produced. IPA isoforms are expressed in approximately 30% of genes across human tissues^7–9^. The expression of IPA isoforms can influence the expression of full-length transcripts. Consequently, IPA dysregulation can alter the levels of full-length proteins, affecting gene functions and contributing to human diseases including cancer^10^. Several studies demonstrated that aberrant IPA transcripts impact cancer development, progression, and therapy resistance^11–14^. We previously showed for the first time that cells from chronic lymphocytic leukemia (CLL) patients generate widespread IPA RNA isoforms^15^. In addition to affecting full-length RNA expression, many IPA-generated truncated proteins exhibit loss of function or dominant-negative effects that functionally disrupt tumor suppressor genes. Of note, we identified IPA expression in tumor suppressor genes in CLL patients who carry no mutation in these genes. This suggested that tumor suppressor genes in CLL could be modulated by either genetic alterations or IPA. Overall, we had proposed IPA-mediated RNA truncation as a mechanism that could result in phenotypic outcomes like those caused by genetic alteration.

In addition to truncated RNA transcripts that lose key protein domain sequences, CLL cells also generate dozens of IPA transcripts that occur in the introns within the 5’ untranslated regions^15^. These IPA transcripts are of particular interest because (1) they do not contain canonical protein sequences of the corresponding parent genes; thus, they were initially categorized as putative non-coding IPA isoforms. (2) These isoforms contain defined polyadenylation signals and polyadenine tails, indicating that they have undergone 3’ end processing. (3) Instead of being recognized as premature products and subjected for degradation, these putative non-coding IPA isoforms have significantly higher expression in CLL cells compared to normal B cells. (4) Many of these isoforms were recurrently found in more than 10% of CLL patients, a patient population with well-known disease heterogeneity^16^. Based on these characteristics, we hypothesized that putative non-coding IPA isoforms are significant IPA changes in CLL cells and they may have pathogenic importance in leukemia.

In this study, we investigated a putative non-coding IPA isoform generated from the *TP53* gene, named *p53-IPA*. The expression of *p53-IPA* was substantially upregulated in leukemia cells in CLL patients compared to their normal counterparts. We verified that *p53-IPA* can produce a protein, IPADP1 (<u>IPA</u>-<u>d</u>erived <u>p</u>rotein 1), from a non-canonical open reading frame. Functional assays revealed that IPADP1 contributes to cancer cell survival, and the survival contribution of IPADP1 was not caused by p53 protein *per se*.

Altogether, we demonstrate that *p53-IPA* is a coding RNA that produces a unique and novel protein from the *TP53* locus. While we and others previously showed that IPA disrupts the parental proteins through IPA-mediated parental gene transcript truncations^9–11,15^, this study demonstrates that IPA isoforms can additionally generate distinct proteins with functions independent of the full-length proteins. Therefore, IPA isoforms have impact beyond regulating or truncating the corresponding parental genes.

## RESULTS

### *p53-IPA* generates a non-canonical protein

We previously identified more than 300 genes that exhibit recurrent IPA events in CLL cells^15^ (events were detected in at least 10% of patients). Among them, 42 genes expressed IPA isoforms in the introns that spanned the 5’ untranslated regions (**Table S1**). To begin elucidating the functional consequences of these putative non-coding IPA isoforms in leukemia cells, we first identified the ones that were generated in genes with reported importance to CLL or other cancers. Among these genes, we investigated an IPA isoform of *TP53*, as it is one of the most important tumor suppressors and has been well characterized in CLL. Moreover, the expression of the IPA isoform produced by the *TP53* gene, referred to as *p53-IPA*, was significantly upregulated in 60% (34 out of 57) of CLL patients ^15^.

*p53-IPA* is 2250 bp long and is generated by the recognition of a polyadenylation site in the 1^st^ intron of *TP53* (**Fig 1A**). As the translation start site of the p53 protein is in exon 2, *p53-IPA* does not include p53 protein sequences (**Fig 1A**). A growing body of experimental evidence has identified unannotated peptides with roles in physiological pathways that can be produced from translatable open reading frames (ORFs) in putative non-coding RNAs^17–19^. This prompted us to determine whether *p53-IPA* could encode a non-canonical ORF. Using ORF Finder (www.ncbi.nlm.nih.gov/orffinder), we found that *p53-IPA* contains two putative ORFs that potentially encode 118 and 139 amino acids, which are predicted to translate into peptides with the sizes of 13 (ORF1) and 15 kDa (ORF2), respectively. To determine if ORF1 and ORF2 were translated from *p53-IPA*, we added a Flag tag to each ORF and ectopically expressed both in HCT116 cells. HCT116 cells are amenable for genetic manipulation and can produce more detectable proteins than leukemia cell lines, which helps validate the translation ability of ORFs. More importantly, HCT116 is a well-characterized colorectal cancer cell line which has wild-type *TP53* and undetectable *p53-IPA* expression. Western blot analysis using an anti-Flag antibody demonstrated that ORF1 can produce a protein in HCT116 cells (**Fig 1B, S1A**), which we named <u>IPA</u>-<u>d</u>erived <u>p</u>rotein 1 (IPADP1). In the Western blot result, the putative ORF2 did not generate protein **Fig S1B**). The DNA and protein sequences of IPADP1 were shown in **Fig 1C**. The protein structure prediction by ColabFold ^20^ showed that IPADP1 has primarily intrinsically disordered regions (**Fig S1C**). Next, we generated a custom-made antibody against the IPADP1 protein. Western blot analysis demonstrated that this antibody recognizes ectopically expressed IPADP1 in HCT116 cells (**Fig 1B, S1A**). The specificity of IPADP1 antibody was further confirmed by reduction of IPADP1 levels in IPADP1-knocked down (KD) cells using shRNA (**Fig S2A, S2B**) and IPADP1-knocked out (KO) cells generated by CRISPR editing (**Fig S2I**). After confirming the specificity and sensitivity of the antibody, we used it to detect the expression of endogenous IPADP1 in 5 blood cancer cell lines, MEC-1 (a CLL cell line), NALM6 (an acute lymphocytic leukemia), K562 (a chronic myeloid leukemia, CML), SC-1 (a B cell lymphoma), and Raji (a Burkitt’s lymphoma). The endogenous IPADP1 expression was found in all 5 blood cancer cells (**Fig 1D, S1D**), whereas the expression was absent in the normal B cell line, BLCL (**Fig 1F**). With cell fractionation to separate cytoplasmic and nuclear fractions of K562 cells, we observed that endogenous IPADP1 expression was enriched in the cytoplasm by Western blot analysis using the anti-IPADP1 antibody (**Fig 1E, S1E**). Consistently, immunofluorescence staining using an anti-Flag antibody showed that ectopically expressed Flag-IPADP1 was enriched in the cytoplasm of HCT116 cells (**Fig S1F**, left). When IPADP1 was fused with an mCherry fluorescent protein in a protein expression plasmid, IPADP1-mCherry protein was also found to be enriched in the cytoplasm of HEK293T cells after plasmid transfection (**Fig S1F**, right). To assess endogenous IPADP1 expression in CLL, we collected 8 samples based on a quality specification that the cryopreserved samples contained >80% tumor cells with at least 90% cell viability after reviving. Western blot analysis using the anti-IPADP1 antibody showed that endogenous IPADP1 expression was detected in 7 out of 8 CLL cells isolated from an independent cohort of CLL patients, with patient #3 (CLL3) exhibiting nondetectable IPADP1 expression (**Fig 1F, S1G**).

**Figure 1.**
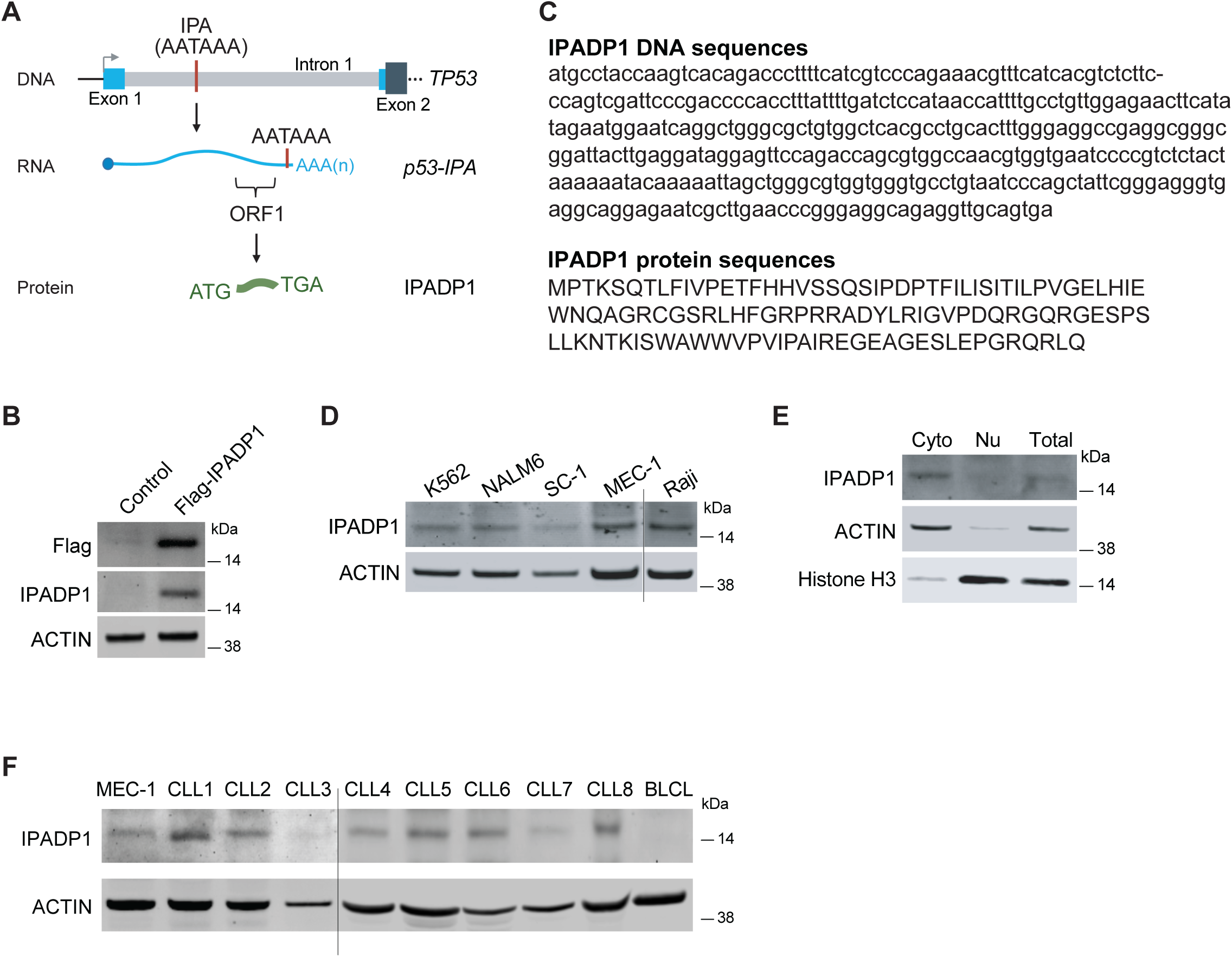
*p53-IPA* generates a non-canonical protein, IPADP1. **(A)** Schematic of the *p53-IPA* isoform. The dark blue box in exon 2 shows the coding region of the p53 protein. The light blue boxes (exon 1 and part of exon 2) are 5’ untranslated regions. **(B)** Western blot analysis of ectopic expression of IPADP1 in HCT116 cells by anti-Flag and anti-IPADP1 antibodies. ACTIN, loading control. **(C)** DNA (top) and amino acid sequences (bottom) of IPADP1. **(D)** Western blot analysis of endogenous IPADP1 expression in blood cancer cell lines with the anti-IPADP1 antibody. ACTIN, loading control. **(E)** IPADP1 expression in K562 with cytoplasmic (Cyto)/nuclear (Nu) fractionation. ACTIN is the control of cytoplasmic fraction and Histone H3 represents the nuclear fraction. **(F)** Endogenous IPADP1 expression was detected in tumor cells derived from CLL patients and a CLL cell line, MEC-1, but not in a normal B cell line, BLCL. ACTIN, loading control.

It is known that in CLL patients, as well as other leukemia patients, *TP53* alterations correlate with more aggressive tumors, lower sensitivity to standard chemotherapies, and shorter survival ^21–23^. Since IPADP1 is a protein derived from the *TP53* locus, we investigated whether IPADP1 affects p53 function. To this end, we first tried to knock down IPADP1 expression by shRNAs in NALM6, a blood cancer cell line with a wild-type *TP53* status. However, the knockdown (KD) of IPADP1 in NALM6 led to cell death shortly after IPADP1 KD, suggesting a survival dependency on IPADP1 in these cells (not shown). The KD control cells (scrambled shRNA and shRNA targeting the luciferase gene) were viable under the same experimental procedures. To help address the question, we used HCT116 cells, as the cells have wild-type *TP53*. Given that p53 function is known to be mainly regulated at the protein level, IPADP1 was ectopically expressed in HCT116 cells to determine whether IPADP1 expression affects endogenous p53 protein levels. IPADP1-expressing HCT116 cells were treated with 5-fluorouracil (5-FU), a chemotherapy agent that can induce p53 accumulation ^24,25^. By Western blot analysis, we found that the expression of IPADP1 did not significantly affect the rate and level of p53 protein accumulation during 5-FU treatments of HCT116 cells (**Fig S1H-I**, with quantified results on the right). The induction of two well-known p53 targets, p21 and MDM2, was also not affected by IPADP1 expression upon 5-FU treatments. The results were consistent in cells treated with either various concentrations of 5-FU (**Fig S1H**) or under various treatment times (6-72 hrs) with 5 µM 5-FU (**Fig S1I**). Next, as the oligomerization of p53 is critical for p53 activity ^26^, we examined whether IPADP1 interferes with p53 oligomerization upon 5-FU treatments. IPADP1-expressing HCT116 cells were treated with 10 µM 5-FU for 16 hrs to induce endogenous p53 accumulation. The cell lysates were treated with glutaraldehyde (GA) ranging from 0.01% to 0.05% (v/v), a protein cross-linking agent that was used to reveal p53 oligomers^27^. As shown in **Figure S1J**, Western blot analysis did not detect significant differences in p53 dimer, trimer, and tetramer formation in IPADP1-expressing HCT116 cells under dosage escalation of GA. Quantified results are shown in **Figure S1J** (on the right, N = 5 biological repeats). This indicates that IPADP1 does not impact p53 oligomerization upon 5-FU treatment. Additionally, we assessed whether IPADP1 could affect p53 function through its association with the p53 protein. We treated IPADP1-expressing HCT116 cells with 5 µM 5-FU for 16 hrs to induce endogenous p53 accumulation and the expression of p21 and MDM2 (as shown in **Fig S1I**). After that, protein pull-down was performed using anti-p53-Trap agarose beads. By Western blot analysis, an interaction of p53 and IPADP1 was not observed under three conditions of p53 pull-down in which the interaction of p53 and MDM2 was detected (**Fig S1K**). Similarly, no significant interaction between p53 and IPADP1 was seen when the protein pull-down assay was performed with anti-Flag-Trap agarose beads (for immunoprecipitation of Flag-IPADP1, data not shown). Collectively, under the conditions tested, IPADP1 did not seem to interact with the p53 protein, nor did it affect p53 protein accumulation and oligomerization.

### The functional impact of IPADP1 on leukemia

Since the IPADP1 expression was observed in CLL patient cells and many blood cancer cell lines, but not normal B cells, we investigated the function of IPADP1 in leukemia cells. We used two independent shRNAs to knock down IPADP1 in MEC-1 and K562 cells, two blood cancer cell lines exhibiting *TP53* mutations or partial deletion of *TP53* ^28,29^. The knockdown efficiency was validated by Western blot analysis for protein expression or qPCR for RNA levels (**Fig S2A, S2B**). We challenged the cells with various chemotherapy agents for blood cancers to evaluate whether IPADP1 can influence the treatment response. With cytotoxicity assays, we found that IPADP1 knockdown in MEC-1 and K562 significantly sensitized the cells to 1 – 40 µM doxorubicin treatment for 24 hrs, a topoisomerase II inhibitor (**Fig 2A, 2B**) ^30,31^. Upon treatments with 0.5 – 5 µM imatinib and 0.25 – 3µM dasatinib for 48 hrs, two CML chemotherapy agents that are frequently used to inhibit the constitutively active tyrosine kinase in patients ^32^, IPADP1 knockdown significantly reduced cell viability in K562 cells (**Fig 2C, 2D**). In both MEC-1 and K562 cell lines, IPADP1-KD resulted in a significant decrease in cell viability during 24 hr treatments with 20 – 80 µM cisplatin, a platinum drug that induces DNA damage and cell death (**Fig S2C, S2D**) ^33^.

**Figure 2.**
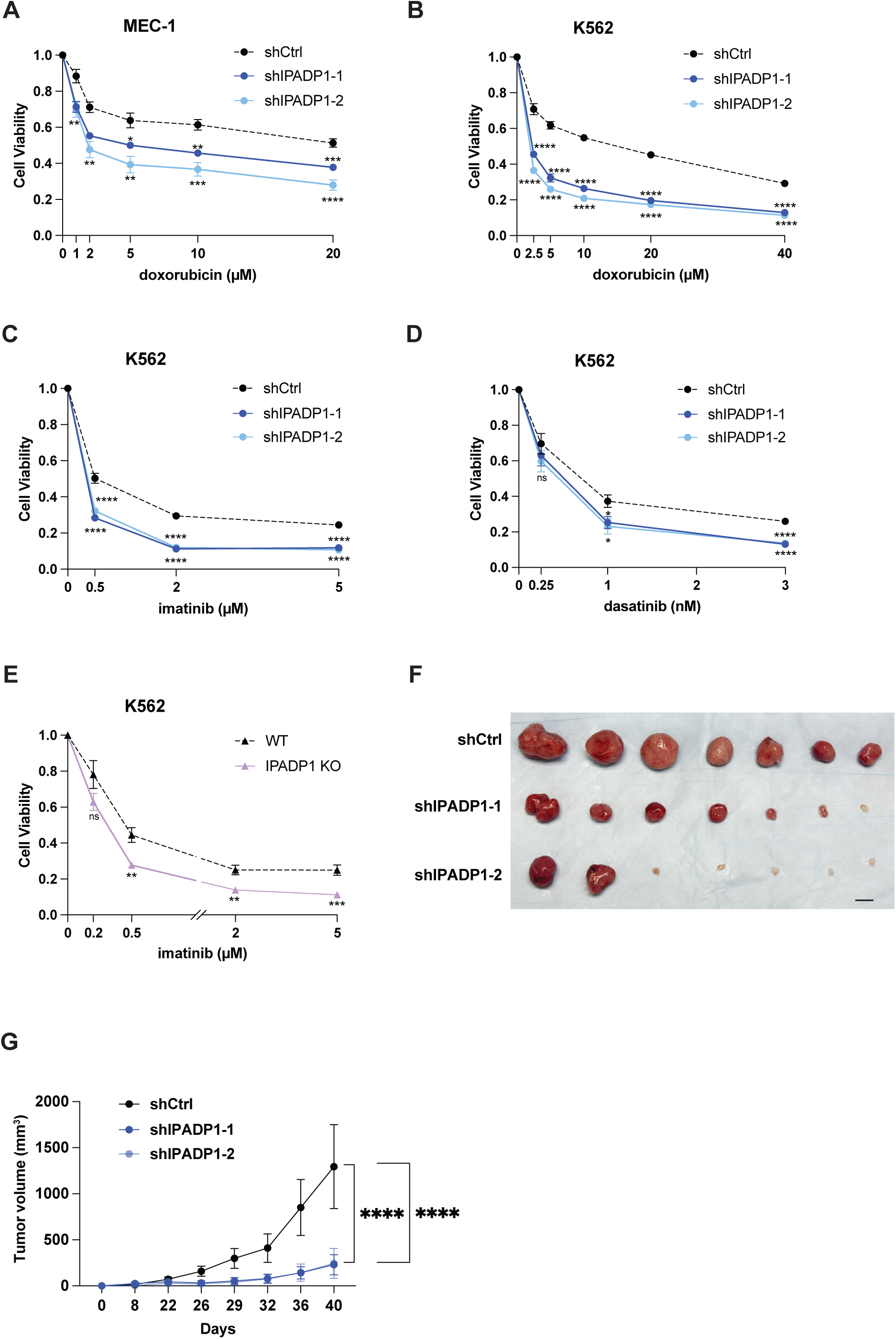
IPADP1 regulates cell viability upon drug treatments as well as xenograft formation *in vivo*. **(A)** Cytotoxicity assays for IPADP1-knockdown (shIPADP1-1,2) and knockdown control (shCtrl) MEC-1 cells upon doxorubicin treatments. Cells were treated with doxorubicin at the indicated concentrations for 24 hrs. Luminescence was detected by the addition of CellTiter Glo. Data was first normalized to a media only control and then to 0 µM within the same cell line. Statistically significant differences were measured by Welch’s t-test unless indicated and are shown by asterisks (*) at a level of *P* < 0.05, (**) at a level of *P* < 0.01, (***) at a level of *P* < 0.001 and (****) at a level of *P* < 0.0001. N = 3 biologically independent experiments. **(B)** As in (A), but performed with K562 cells treated with doxorubicin for 24 hrs. N = 3 biologically independent experiments. **(C)** As in (A), but performed with K562 cells treated with imatinib for 48 hrs. N = 3 biologically independent experiments. **(D)** As in (A), but performed with K562 cells treated with dasatinib for 48 hrs. N = 3 biologically independent experiments. **(E)** As in (A), but performed with K562 wild-type (WT) and IPADP1-CRISPR-knockout (IPADP1-KO) cells treated with imatinib for 48 hrs. N = 3 biologically independent experiments. **(F)** Xenograft formation assay in nude mice with K562 control (shCtrl) and two IPADP1-knockdown (shIPADP1-1,2) cells. Scale bar = 1 cm. **(G)** The measurement of tumor size from at least 7 mice in each group. \*\*\*\**P* < 0.001

We also performed cytotoxicity assays to assess the role of IPADP1 ectopic expression in HCT116 cells. Consistently, IPADP1 expression resulted in enhanced cell survival upon the 72 hr treatment with 2.5 – 10 µM 5-FU in HCT116 cells (**Fig S2E**). In addition to cytotoxicity assays, clonogenic assays were performed to further evaluate the impact of IPADP1 on the growth and survival of IPADP1-expressing HCT116 cells with 5-FU treatment. Our results showed that treated IPADP1-expressing cells were able to form more colonies compared to treated control cells (**Fig S2F**). Together, these results indicate that IPADP1 contributes to survival advantages in tumor cell lines upon drug treatments.

To further verify the role of IPADP1 in regulating cell survival upon drug treatments, we mutated the translation start site of IPADP1 by CRISPR editing (IPADP1 knockout or IPADP1 KO) in K562 cells. Note that this approach could not be performed in MEC-1 cells due to massive cell death caused by electroporation of preformed Cas9 and guide RNA ribonucleoprotein. The mutations in K562 were confirmed by Sanger DNA sequencing (**Fig S2G**). Consequently, the mutated cells express *p53-IPA* RNA but cannot produce IPADP1 protein. Cytotoxicity assays showed that IPADP1-KO cells exhibit lower viability during the treatments of imatinib and doxorubicin (**Fig 2E, S2H**). The decrease of cell viability in the treated IPADP1-KO cells was similar to the effect seen in treated IPADP1-KD cells (**Fig 2C** and **2E**). This demonstrates that the survival advantage in cells could be conferred primarily by the IPADP1 protein.

Next, we investigated how cell viability could be restored by re-introducing IPADP1 expression in the IPADP1-KO cells. By Western blot analysis using the custom-made anti-IPADP1 antibody, we detected IPADP1 expression in IPADP1-KO cells with the ectopic expression of full-length *p53-IPA* (**Fig S2I**). This shows that in addition to HCT116 cells, ORF1 within *p53-IPA* is translatable in K562 cells. Performing cytotoxicity assays, we found that the rescue of IPADP1 expression could significantly improve, although not fully recover, cell survival in imatinib treated IPADP1-KO cells (**Fig S2J**).

Prompted by the observations that IPADP1 plays a role in regulating cell viability in leukemia cell lines, we next evaluated the oncogenic properties of IPADP1 *in vivo*. One million of the IPADP1-KD or control K562 cells were subcutaneously implanted in the immunodeficient nude mouse (7 mice per group). The mice were monitored for tumor formation for 40 days until the volume of tumors formed by control K562 cells reached to 1500 mm^3^. Our results showed that KD of IPADP1 significantly abolished tumor formation and growth in mice, resulting in a strikingly lower incidence of xenograft formation (**Fig 2F, 2G**). Most xenografts formed by control K562 cells had sizes larger than a diameter of 1 cm (**Fig 2F**, scale bar = 1 cm). On the contrary, mice inoculated with IPADP1-KD cells had either small or no detectable tumor. The body weight of mice was not significantly affected (**Fig S2K**). These results clearly indicated that IPADP1 contributes to tumor formation *in vivo*.

### IPADP1 interacts with multiple proteins in survival related pathways

To elucidate mechanistically how IPADP1 promotes tumor formation *in vivo* and regulates apoptotic response in cell lines during drug treatments, we leveraged proteomic tools to identify proteins that interact with IPADP1. To this end, either a Flag-IPADP1 or control plasmid was expressed in HEK293T (293T) cells, as 293T cells produce sufficient amount of IPADP1 for protein immunoprecipitation (IP) and proteomic analysis. Also, IPADP1 expression is enriched in the cytoplasm of HEK293T cells (**Fig S1C**), similar to leukemia cell line (**Fig 1E**). Coupling protein IP using anti-Flag-Trap agarose beads with mass spectrometry analysis, the result revealed hundreds of potential interactors of IPADP1. While the number of interactors may seem unexpectedly high based on the relatively small size of IPADP1, the intrinsically disordered regions of IPADP1 may contribute to its structural flexibility and ability to interact with diverse proteins.

Gene Ontology analysis of mass spectrometry data revealed the enrichment of IPADP1 interactome proteins in several processes that are critical for cell survival, such as cell cycle, apoptosis, DNA repair, and DNA replication (**Fig 3A**, **Table S2**). Based on our data that IPADP1 localizes mostly in the cytoplasm and regulates cell viability, we validated by Western blot analysis the interactions of IPADP1 with cell death-related proteins BAG6, AIFM1, BAX, DNM1L and cell cycle protein CDK1 which is also a E2F target, as well as DNA repair related protein MSH2 and UMPS in 293T cells (**Fig S3A**). The interaction of IPADP1 to SRP54 was non-significant (**Fig S3B**). Furthermore, we validated the interactions of IPADP1 with all interactors in K562 cells. The interactions of IPADP1 to BAG6, AIFM1, BAX, DNM1L, CDK1 and MSH2 were faithfully observed (**Fig 3B**). In both 293T and K562 cells, the interaction of IPADP1 to UMPS was less enriched than other interactors (**Fig S3A, S3C**).

**Figure 3.**
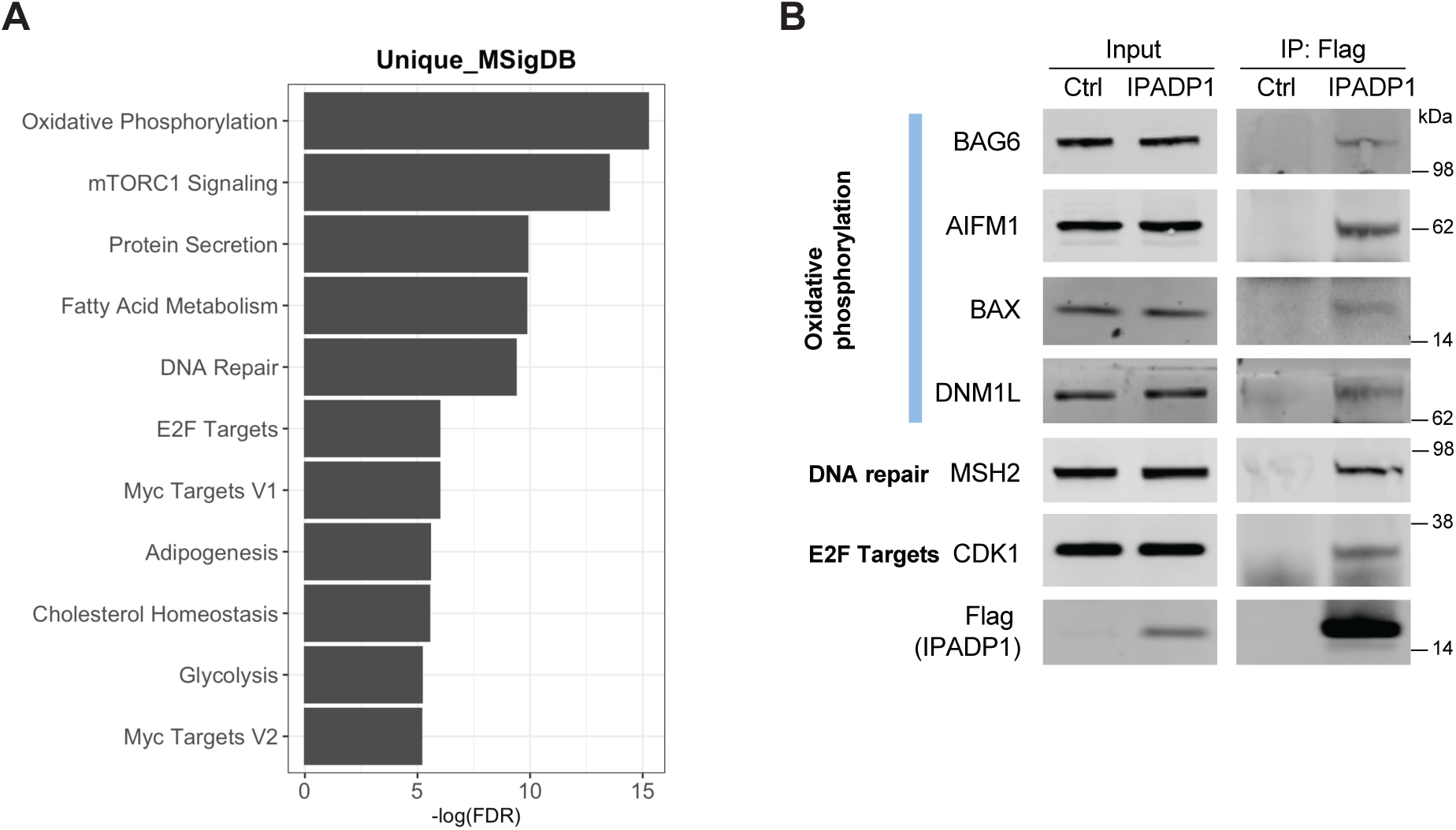
The identification and validation of interactors of IPADP1. **(A)** Gene Ontology analysis of mass spectrometry results for IPADP1 interactors. **(B)** Western blot validation of IPADP1 interactors in K562 cells. The interactors were identified by Flag protein immunoprecipitation in the control or IPADP1-expressed cells. 0.5% input was loaded.

### IPADP1 inhibits apoptosis during drug treatments

Our proteomic analysis results showed that IPADP1 binds to proteins involved in the regulation of cell death, suggesting that IPADP1 may promote cell viability and tumor formation through regulation of apoptosis. To test the hypothesis, we performed flow cytometry analysis with the staining of annexin V, an early apoptosis marker, and DAPI, a DNA-binding dye for DNA content in cells, to examine how IPADP1 affects cell death upon drug treatments ^34^. DAPI and annexin V double-positive staining indicates cells in the late stage of apoptosis^35^.

We treated the control and IPADP1-knockdown (KD) K562 cells with low concentrations of drugs based on the results of cytotoxicity assays (**Fig 2B-2D, S2D**) to induce apoptosis in cells. We found that KD of IPADP1 led to a significant 3 to almost 5-fold increase in the percentage of annexin V and DAPI double-positive K562 cells upon doxorubicin treatments (**Fig 4A**), 2 to 3.5-fold increase upon cisplatin treatments (**Fig 4B**) and 5-fold increase with imatinib treatments (**Fig 4C**). The results were consistent between two IPADP1-KD cell lines. Moreover, annexin V single-positive cells were consistently enriched in IPADP1-KD cells during the treatments with all three drugs, further supporting the idea that IPADP1 may prevent cells from entering apoptosis processes (**Fig S4A-S4C**). A similar increase in the apoptotic population was also seen when the IPADP1-KD MEC-1 cells were treated with doxorubicin (**Fig S4D**). Together, the results suggest that IPADP1 has an anti-apoptotic role upon drug treatments in blood cancer cells.

**Figure 4.**
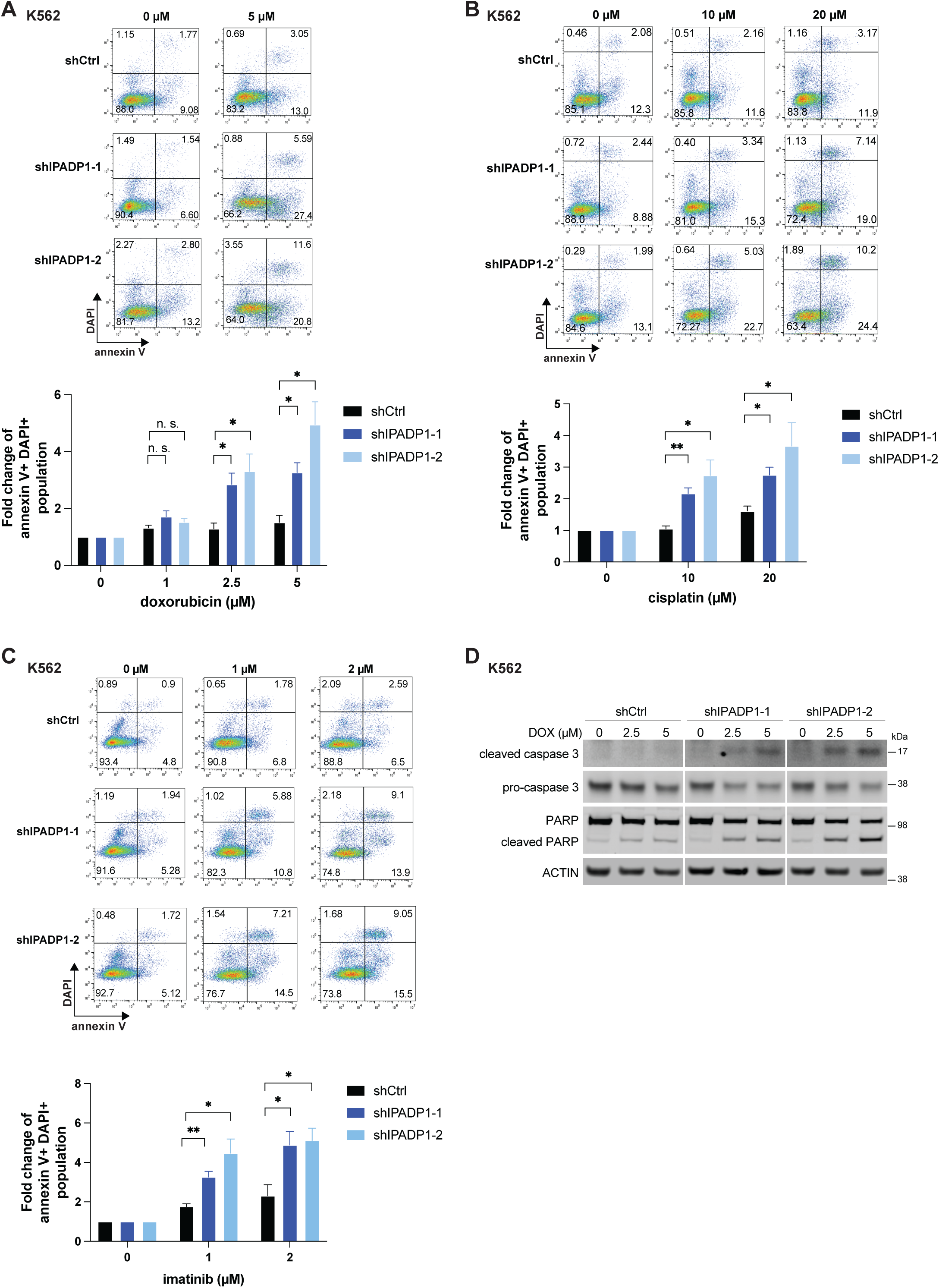
IPADP1 inhibits apoptosis upon drug treatments. **(A)** Top, the flow cytometry analysis for apoptosis in control and IPADP1 knockdown K562 cells treated with doxorubicin. Cells were treated with 0 µM and 5 µM doxorubicin for 24 hrs and stained with annexin V and DAPI. Unstained and single stained controls were used to assess compensation. Bottom, the result from independent experiments (N = 3). \**P* < 0.05, n. s., non-significant. **(B)** As in (A), but performed in cells treated with 0 µM, 10 µM, and 20 µM cisplatin for 24 hrs. N = 3 biologically independent experiments. \**P* < 0.05, \*\**P* < 0.01. **(C)** As in (A), but performed in cells treated with 0 µM, 1 µM, and 2 µM imatinib for 48 hrs. N = 3 biologically independent experiments. \**P* < 0.05, \*\**P* < 0.01. **(D)** Western blot analysis of K562 control and shRNA-mediated IPADP1 knockdown cells treated with indicated concentrations of doxorubicin (DOX) for 16 hrs. ACTIN, loading control.

Additionally, Western blot analysis confirmed that doxorubicin treatments of IPADP1-KD K562 cells led to the expression of cleaved caspase 3, an indicator of apoptotic response. On the contrary, the cleaved caspase 3 expression was absent in control cells under the same treatment conditions (**Fig 4D**). This shows that the anti-apoptotic effect of IPADP1 is caspase-dependent. Consistently, treated IPADP1-KD cells exhibit higher levels of cleaved PARP, a cellular hallmark of caspase-mediated apoptosis ^36^.

### IPADP1 regulates DNA damage response, DNA repair, and cell cycle

We identified IPADP1 associated interactions with DNA repair related proteins, such as RAD21, RAD23B, SMC1, UMPS, and MSH family genes MSH2, MSH4, and MSH6 ^37,38^ and validated the interaction of IPADP1 to MSH2 and UMPS, leading us to hypothesize that IPADP1 may regulate DNA damage response and repair, in addition to apoptosis (above).

Doxorubicin is a Topoisomerase II inhibitor and causes DNA double-strand breaks (DSBs) ^39^. When DSBs occur, H2AX is rapidly phosphorylated to form γ-H2AX, marking the damage sites and recruiting DNA repair proteins ^40,41^. Therefore, γ-H2AX signal is used for DSB detection and the evaluation of cellular responses to DNA damage ^42,43^. To investigate if IPADP1 impacts cellular DNA damage response, the IPADP1-KD and control K562 cells were treated with doxorubicin 5 µM for one hour and then recovered at various timepoints (3 to 48 hrs) after removing the drug. Following γ-H2AX staining, flow cytometry analysis was applied to examine the DNA damage response (DDR) and repair ability of cells.

We found that after 1-hr doxorubicin treatment, the γ-H2AX signal in IPADP1-KD cells was approximately half of that in control cells (**Fig 5A**, 0 hr recovery). At 3 hrs after drug removal, the γ-H2AX signal in IPADP1-KD cells remained ∼33% less than that in control cells. This suggested that IPADP1 may regulate the phosphorylation of H2AX. Later, at 6 hrs after drug removal, the γ-H2AX signal decreased significantly in the control cells, indicating the repair of DNA damage by DDR. On the contrary, the γ-H2AX signal in IPADP1-KD cells remained unchanged even at 48 hrs after drug removal. The retention of DNA damage indicates DDR deficiency in IPADP1-KD cells. The results together suggest that IPADP1 may play a role in DNA damage response in cells.

**Figure 5.**
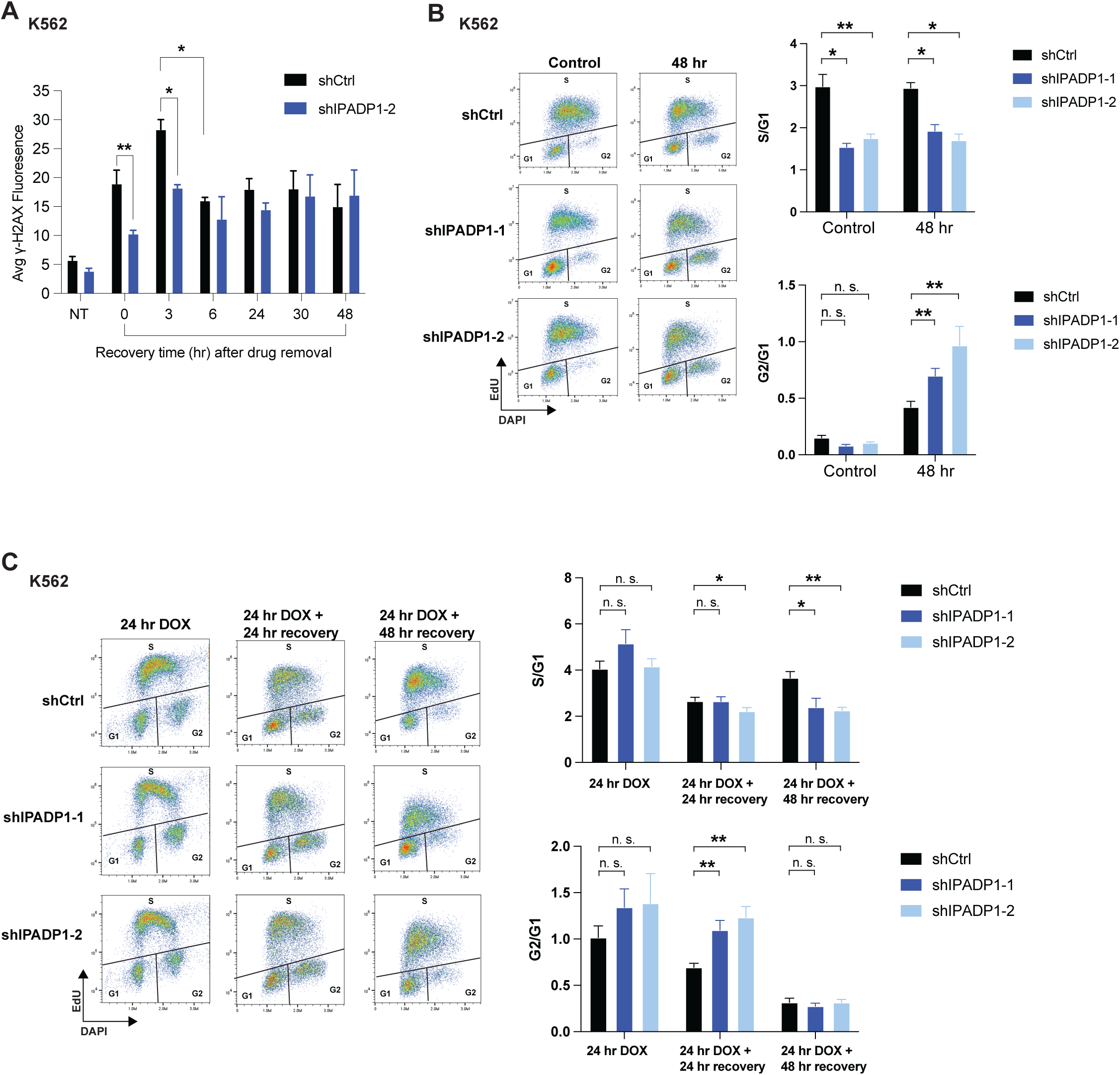
IPADP1 regulates DNA damage response, repair, and cell cycle. **(A)** The flow cytometry analysis of γ-H2AX signal in K562 shRNA-mediated IPADP1 knockdown and control cells. Cells were treated with 5 µM doxorubicin for 1 hr. After that, the drug was removed, and cells were recovered for the indicated timepoints. At each timepoint, cells were harvested and subjected for γ-H2AX staining and flow cytometry analysis. Statistical differences shown are between shCtrl and shIPADP1-2 at 0 and 3 hr recovery as well as between shCtrl cells at 3 and 6 hr recovery. The difference between 3 and 6 hr recovery in shIPADP1-2 knockdown cells was not significant. N = 4 biologically independent experiments. \**P* < 0.05, \*\**P* < 0.01. **(B)** Flow cytometry analysis of cell cycle progression in K562 cells. Control and IPADP1 knockdown cells were treated with 20 nM doxorubicin or DMSO for 48 hrs, followed by EdU and DAPI staining (left). Quantification of flow cytometry results shown on the right. N = 3 biologically independent experiments. \**P* < 0.05, \*\**P* < 0.01, n. s., non-significant. **(C)** As in (B) but done with cells treated with 20 nM doxorubicin for 24 hrs plus 24 hr and 48 hr recovery. Quantification of flow cytometry results shown on the right. N = 3 biologically independent experiments. \**P* < 0.05, \*\**P* < 0.01, n. s., non-significant.

In addition to DNA damage response proteins, mass spectrometry showed that IPADP1 interacts with proteins that are involved in the control of the cell cycle, such as CDK1, a cyclin-dependent kinase, and CCNA2, a cyclin protein ^44,45^.

To investigate the role of IPADP1 in cell cycle regulation, EdU incorporation was used to assess for DNA synthesis (cell proliferation), and DAPI staining was used to show DNA content which helps identify cells at G1 and G2 phases. With flow cytometry analysis, we found that IPADP1-KD K562 cells exhibit less EdU signals (less DNA synthesis events) and more G0/G1 arrest than control cells, showing as >40% lower S/G1 in **Fig 5B** (Control, the top bar graph). This suggested that IPADP1 may play a role in G1-to-S transition. Doxorubicin can cause cell cycle arrest at G1/S and G2/M stages ^46^. We treated control and IPADP1-KD K562 cells with a low dose of doxorubicin at 20 nM for 48 hrs, a treatment condition which did not cause drastic overall cell death but can induce cell cycle changes. The flow analysis results showed that upon the treatment, IPADP1-KD cells were less EdU-positive and more G2 arrested than control cells (**Fig 5B**, 48 hr). This suggests that either IPADP1 could regulate cell cycle steps beyond S phase, or the defect caused by IPADP1 knockdown during S phase could continue to impact G2 progression in IPADP1-KD cells. Using the same assay for HCT116 cells, we found that, consistently, the ectopic expression of IPADP1 in cells with p53-null background significantly enhanced DNA synthesis upon 5 µM 5-FU treatment for 72 hrs, with less cells retained at G2 (**Fig S5A**). Together, these results suggest that IPADP1 may promote cell transition to S phase and decrease G2 arrest during drug treatments. The data also provides an insight into how IPADP1 may facilitate tumor formation and cell survival through regulating cell cycle progression.

DNA repair defects can cause cell cycle arrest ^47^. Based on our observations thus far, we next investigated whether IPADP1 prevents cells from arresting at the G2 phase and facilitates cell cycle progression by regulating cell recovery from drug treatments. The control and IPADP1-KD K562 cells were subjected to 20 nM doxorubicin for 24 hrs and then had either 24 or 48 hrs recovery after drug removal. The same EdU and DAPI staining assay was performed to examine cell cycle changes. The 24-hr doxorubicin treatment did not significantly alter cell cycle progression in control and IPADP1-KD cells. Later, upon 24 hrs recovery after drug removal, we observed that more IPADP1-KD cells were retained at G2 compared to control cells (**Fig 5C**, the middle panel in the bottom bar graph). This suggests that IPADP1 may influence how cells recover from doxorubicin-induced cell cycle arrest. After 48 hr recovery, all cells showed reduced G2 retention, suggesting that cells may exit G2 arrest. However, compared to control cells, both IPADP1-KD cells showed less EdU staining, suggesting that cells lacking IPADP1 may not be able to enter DNA synthesis and were retained at G0/G1. This finding indicates that IPADP1-KD cells have defective cell cycle progression under the recovery conditions. Overall, our data suggest that IPADP1 may play a role in the recovery of doxorubicin-induced cell cycle arrest.

### *p53-IPA* expression is associated with *SF3B1* mutations

We previously reported that IPA events are more frequently found in CLL patients with mutations in *SF3B1*, a core factor in the U2 spliceosome^48^, than the individuals without the mutations^15^. To assess whether the expression of *p53-IPA* could be associated with *SF3B1* mutations, we investigated an independent cohort of 29 CLL patients whose *SF3B1* mutation status was known. 10 out of 29 patients carry mutations of *SF3B1*. The expression of *p53-IPA* was examined using QuantSeq, a commercialized 3’-based sequencing method, and APAlyzer was performed to analyze the results^49^. The expression of *p53-IPA* was found in 6 out of 29 patients (**Table S3**). Importantly, we observed that the expression of *p53-IPA* is significantly associated with *SF3B1* mutations (Odd ratio = 5.67, *p* = 0.046, **Fig 6A**). In addition to CLL, *SF3B1* mutations are commonly found in myelodysplastic syndromes (MDS), a group of blood cancers characterized by defects in blood cell maturation^50,51^. We examined *p53-IPA* expression in a cohort of 65 MDS patients with known *SF3B1* mutation status using QuantSeq. *p53-IPA* upregulation was observed in 10 out of 65 MDS patients and the *p53-IPA* expression was also more prevalent in MDS patients bearing *SF3B1* mutations than wild-type individuals (Odd ratio = 4.46, *p* = 0.03, **Fig 6B**). To support the correlation, we examined *p53-IPA* expression in isogenic K562 cells lines with wild-type or mutated *SF3B1*. qPCR results showed that *p53-IPA* expression is significantly higher in mutated cells (**Fig 6C**), suggesting that *SF3B1* mutations may impact the expression of *p53-IPA* in hematologic malignancies.

**Figure 6.**
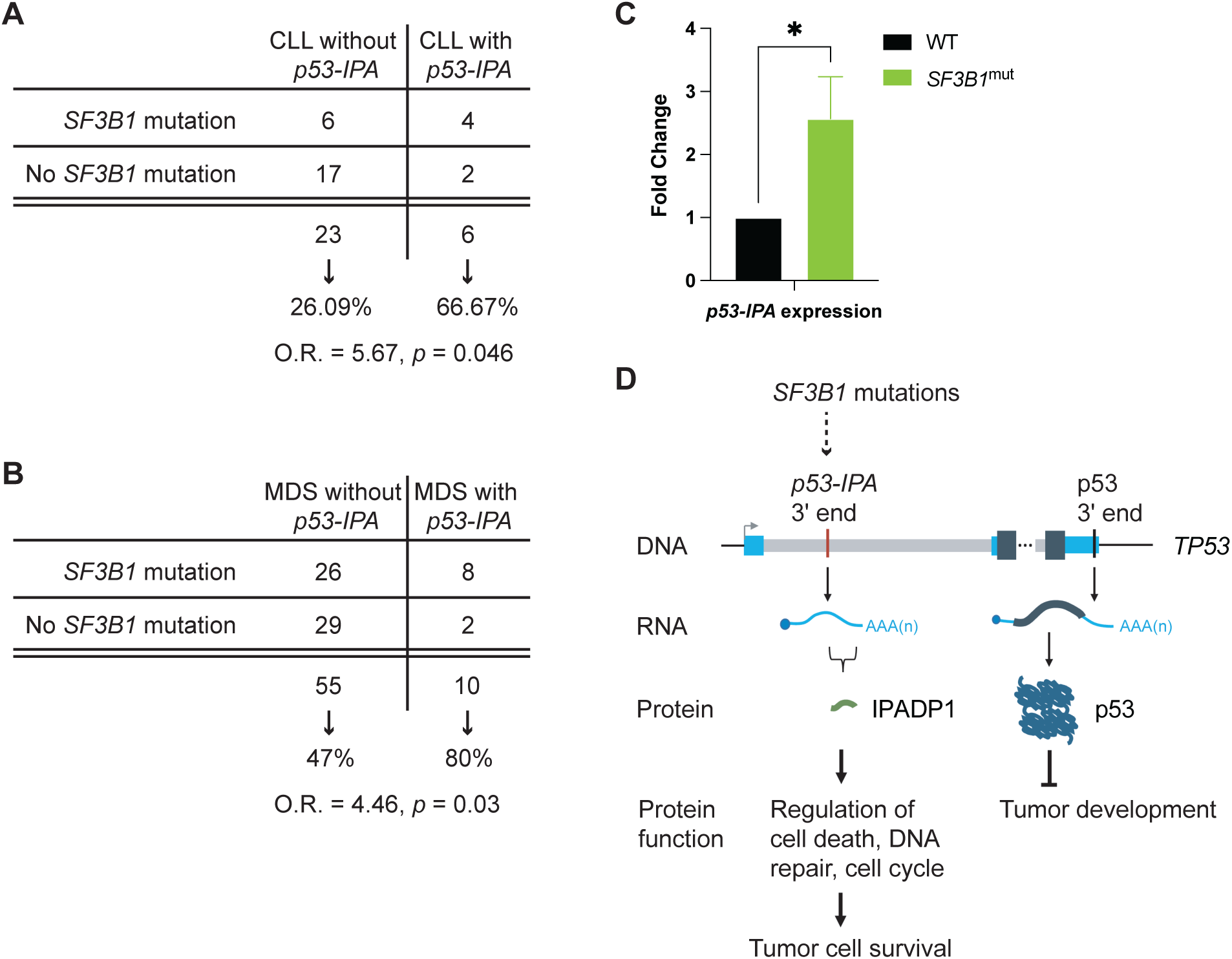
*p53-IPA* expression is more prevalent in patients with *SF3B1* mutations than patients without the mutations. **(A)** *p53-IPA* expression is significantly associated with *SF3B1* mutations in CLL. O. R., odd ratio. Statistical significance was assessed using Boschloo’s exact test conditioned on the fixed *p53-IPA* positive margin. **(B)** As in (A), the association of *p53-IPA* expression and *SF3B1* mutations is significant in MDS patients. **(C)** qRT-PCR result of *p53-IPA* expression in wild-type (WT) or *SF3B1* mutated (*SF3B1*^mut^) K562 cells. N = 6 biologically independent experiments. \**P* < 0.05. **(D)** IPADP1 has functions independent of its parental gene. IPADP1 promotes cell survival, whereas p53 is a well-recognized tumor suppressor.

## DISCUSSION

### IPA isoforms can produce a functional non-canonical protein

In this study, we functionally characterized a *TP53* IPA isoform (*p53-IPA*) in patient leukemia cells. The expression of *p53-IPA* was recurrent and significantly higher in many patient’s cells than in normal B cells^16^. We showed that *p53-IPA* encodes a previously unidentified protein, IPADP1, that does not overlap p53 protein sequences.

IPADP1 and p53 protein are produced from the same locus. Under the conditions that we tested to activate p53 protein, IPADP1 did not disrupt p53 protein accumulation or oligomerization nor did it interact with p53. This suggests that IPADP1’s role in leukemia cells may not be through impacting p53 protein function. Performing cell-based and xenograft assays, we found that IPADP1 has tumor-promoting properties (Summary in **Fig 6D**). Given that full-length p53 protein is well-recognized to be tumor-suppressive, our data indicates that IPA isoforms can exhibit properties different from their full-length products, and genes can generate products with distinct functions through IPA. In the example of IPADP1 and p53, these two products do not share the same functional domains. Consequently, IPA helps diversify gene functions beyond coding sequences, as well as expand gene products.

### IPADP1 is intrinsically disordered with poor conservation

Evolutionary conservation can be used as an indicator of functionality prediction. However, since IPADP1 is intron-derived, the sequences do not show significant conservation between human and most organisms based on the BLAST results of DNA and protein sequences. While several amino acids in a short region of IPADP1 showed partial similarity with primate proteins, the proteins and their domain functions were hypothetical (data not shown). Similarly, no functional protein domain was predicted using either CD-Search (https://www.ncbi.nlm.nih.gov/Structure/cdd/wrpsb.cgi), InterPro (https://www.ebi.ac.uk/interpro/), or PROSITE (https://prosite.expasy.org/). Structurally, Colabfold^20^ showed that IPADP1 is mostly intrinsically disordered, with only one condition exhibiting an acceptable confidence (pLDDT score 70-80) to predict three putative short beta-sheet regions (**Fig S1C**). It has been shown that intrinsically disordered proteins could have high binding capacity because their unstructured domains provide flexibility such as folding-upon-binding and conformational buffering to adopt the structures of their binding partners^52^. This could contribute to their diverse specificity to the binding proteins, as well as their multifunctionality in biological processes. Consistent with the observations, our proteomic analysis identified hundreds of IPADP1 interactor candidates. Moreover, Western blot analysis validated 7 out of 8 interactors of IPADP1 in both 293T cells and K562 cells. This suggested that a proportion of proteins identified by mass spectrometry analysis may be true IPADP1 interactors. Therefore, IPADP1 could be a hub of protein networks that engage in many fundamental pathways. Supporting this idea, the validated interactions of IPADP1 with BAX, BAG6, DNMT1, MSH2, and CDK1 help uncover IPADP1’s roles in regulating cell death, DNA damage response, and cell cycle progression.

### IPADP1 may have cancer-promoting contributions beyond CLL

In addition to the CLL cell line MEC-1, the survival advantage of IPADP1 expression was also found in K562, a CML cell line. Moreover, IPADP1 expression was detected in acute leukemia and lymphoma cell lines. This indicates that IPADP1 may have a cancer-promoting impact across blood cancers. Furthermore, ectopic expression of IPADP1 enhanced HCT116 cell viability during 5-FU treatments, suggesting that IPADP1 contributes to cell survival beyond blood cancers. Supporting this hypothesis, besides CLL, *p53-IPA* expression was detected in MDS patients (this study), as well as in solid tumors such as bladder and ovarian cancers, although the frequency of *p53-IPA* occurrence among patients with solid tumors was unclear ^53,54^. Of note, a recent study reported that *p53-IPA* contributed to melanocyte formation in a zebrafish melanoma model (*p53^-/-^;mitfa:BRAF^V600E^;mitfa^-/-^*) ^54^. Overall, these findings suggest that IPADP1 may play roles in many cancers other than CLL.

### The underlying mechanisms of *p53-IPA* formation

The molecular mechanisms by which *p53-IPA* occurs in CLL remain elusive. We observed *p53-IPA* and IPADP1 expression in treatment-naïve CLL patients and blood cancer cell lines, suggesting that the expression could be induced during tumor development/progression without drug treatments. We further observed that *p53-IPA* expression is significantly associated with *SF3B1* mutations in CLL and MDS and the expression is upregulated in leukemia cell line K562 bearing *SF3B1* mutations. This data suggests an important mechanistic link that *SF3B1* may play a role in regulating *p53-IPA* expression. *SF3B1* is the most commonly mutated splicing factor in CLL. CLL patients with *SF3B1* mutations show poor survival and treatment response. As *p53-IPA* has oncogenic properties, understanding the causal link of *SF3B1* mutations and *p53-IPA* expression may help improve understanding of CLL survival pathways and may suggest treatment options. Some of CLL and MDS patients in the cohorts with known *SF3B1* mutation status had received treatments. We do not rule out the possibility that *p53-IPA* expression in some treated patients could be treatment-related, in addition to *SF3B1* mutations. Future studies including longitudinal specimens will help delineate the relationship of *SF3B1* mutations and *p53-IPA* expression.

Patients without *SF3B1* mutations also express *p53-IPA*, suggesting that *p53-IPA* expression can be regulated by lesions other than *SF3B1* mutations. The genesis of IPA suggests that the typical splice sites for intron excision might be poorly used, allowing to the usage of IPA sites instead. This could be due to defects in factors that recognize splicing motifs. During RNA processing, the U1 spliceosome primarily recognizes 5’ splice sites. Mutations in U1 spliceosomal RNAs are found in approximately 3.5% of CLL patients and 10-30% in B cell lymphomas ^55,56^. However, thus far, the causality of U1 mutations and IPA regulation has yet to be determined. Besides U1, a recent preprint demonstrated that the dysregulation (knockdown, inhibition) of key splicing factors induced IPA expression ^57^. Similarly, numerous studies have also shown that alternative RNA isoforms can be generated through the altered coordination between RNA processing factors ^58,59^. Taken together, kinetic changes between these proteins may favor the recognition of polyadenylation signals in introns over splice sites, resulting in *p53-IPA*. To date, a great deal of research has shown that patients with hematological disorders frequently bear genetic alterations in RNA processing factors and RNA binding proteins ^51,60,61^, suggesting that some alterations, if not all, may result in the occurrence of CLL-IPA.

A recent study revealed that 2 out of 485 CLL patients carried mutations at the 5’ splice site of the intron 1 of *TP53* ^62^. While this affirms the low mutational burden observed in CLL, this particular type of mutation results in the loss of a canonical 5’ splice site, which may lead to alternative splicing or the retention of intron 1. This could contribute to the expression of *p53-IPA* and IPADP1. To date, no mutations have been reported for the polyadenylation signal of *p53-IPA*.

It has been shown that IPA transcripts could result from defects in RNA degradation-associated factors, such as CDK13 and the PAXT complex ^54,63^. Moreover, the loss of a transcription-associated kinase CDK12, either by genetic mutations or CDK12 inhibitor, leads to the formation of IPA in DNA damage response genes in ovarian and prostate cancers, as well as neuroblastoma ^64,65^. However, no DNA lesions in *CDK12*, *CDK13*, or components of the PAXT complex have been reported in CLL. None of these genes located in the chromosome regions that are commonly deleted in either CLL or other leukemia. Future investigations would be needed to determine the molecular basis of *p53-IPA* expression in blood cancers.

### IPA-derived functional non-canonical proteins can be widespread

It is known that RNA 3’ end processing helps stabilize RNA and regulates RNA transport and translation^66^. Our sequencing method identified RNA isoforms with polyadenylated tails and genuine polyadenylation signals in cells. As the discovery of new transcripts outside of annotated protein regions remains challenging, our strategies present a new framework to uncover functional RNA isoforms in cells.

The vast majority of the human genome currently remains annotated as non-coding. Including our studies, an increasing body of work has been devoted to demonstrating that unannotated proteins can be produced from the preconceived non-coding regions ^19,67^. Beside introns, researchers have proposed that many untranslated regions and pseudogenes likely have translation potential ^68^. To test this hypothesis, investigations employing translation and proteomics analysis such as ribosome profiling and mass spectrometry will provide insights into the translational output of these regions, including IPA isoforms, and help illuminate how these non-canonical coding regions impact the proteome. These techniques, combined with the methods of investigation used in this study, will provide powerful tools to reveal the importance of the largely uncharacterized regions of the genome.

Overall, our work underscores the important contribution of IPA isoforms to cancer pathogenesis. It highlights that IPA could be an important “non-genetic” cancer-driving mechanism, and its role might be even more significant in tumors that carry a low mutational burden such as liquid tumors. Our findings have great potential to discover new therapeutic targets and suggest treatment strategies for these patients.

## Acknowledgement

We thank Swathi Arur, Richard Behringer and Kuo-Shun Hsu for their constructive comments and insights into the manuscript. We also thank Prathyushasai (Sai) Machineni, Brian Nyakiti, Van Chanh “Quy” Le and Michael Yao for their technical support for some assays in this study. We thank Chun-Wei (Alan) Chen for helping with the gene ontology analysis of IPADP1 interactors. We thank Nicole Vaughn and the Flow Cytometry and Cellular Imaging Core Facility at MD Anderson for flow cytometry cell sorting and analysis. We thank the Taplin Mass Spectrometry Facility at Harvard Medical School for proteomic studies. We appreciate Rong Yao, Eric Sisson, Stan Bujnowski for providing technical support for high-performance cluster (HPC) resource (http://hpcweb.mdanderson.edu/citing.html). Cell authentication was performed by the Cytogenetics and Cell Authentication Core at MD Anderson Cancer Center.

This study was supported by the following grants: CPRIT (#RR210070 to L.A.C., K.L.L., C.A.O., S.M.F., S.K., X.H.), UT STARs Program and MD Anderson StartUp Fund (S.-H.L.), American Cancer Society Postdoctoral Fellowship PF-25-1433660-01-PFCBI (Y.-W.H.), NIH/NCI R01CA246130 (R.Z. and D.-F.L.), DoD RCRP HT9425-24-1-0957 and DoD RCRP HT9425-25-1-0893 (D.-F.L.), NIH/NIDDK 5U01DK131383 (A.H.T.), Institutional Research Grant from MD Anderson Cancer Center and NIH R01HL173597 (M.J.Y.). D.-F.L. and S.-H.L. are CPRIT Scholars in Cancer Research. S.X.L. is a C. R. Krishnamurthi Faculty Scholar and is supported by awards from the Gabrielle’s Angel Foundation, the Doris Duke Charitable Foundation and the MDS Foundation. M.S. is supported by the Gibson Family Foundation. The Flow Cytometry and Cellular Imaging Core Facility at MD Anderson was supported in part by The University of Texas MD Anderson Cancer Center, P30CA016672, and a Shared Instrumentation Award from the Cancer Prevention Research Institution of Texas (CPRIT).

## Author Contribution

Conceptualization: S.-H.L. Resources & Methodology: R.Z., D.-F.L., A.H.T., S.X.L., M.J.Y., S.-H.L. Investigation: L.A.C., K.L.L., C.A.O., S.M.F., S.K., X.H., Y.-W.H., A.S., M.S., F.Y. Funding acquisition: R.Z., D.-F.L., A.H.T., S.X.L., M.J.Y., S.-H.L. Project administration: S.-H.L. Supervision: R.Z., D.-F.L., A.H.T., S.X.L., M.J.Y., S.-H.L. Writing: L.A.C., K.L.L., S.M.F., S.-H.L., with the support and input from all authors.

## METHODS

### EXPERIMENTAL MODEL AND STUDY PARTICIPANT DETAILS

#### Patient Samples

MD Anderson Cancer Center cohort-Samples were obtained from untreated CLL patients seen at MD Anderson Cancer Center. All patients were consented based on IRB protocol #LAB10-0682. Only de-identified specimens were used in this study. Peripheral blood mononuclear cells from CLL samples with a minimum white blood cell count of 18,900 per microliter were isolated by Ficoll (Cytiva, USA, 17144002) gradient centrifugation at 1200 r.c.f. for 10 min, followed by two washes in PBS at room temperature. Cells were treated with red blood cell lysis buffer for 5 min at room temperature and were washed once with PBS. After that, cell numbers were determined before storage at -80°C or liquid nitrogen. All CLL samples in our study had tumor cells making up more than 80% of the blood cells.

Stanford University cohort-Healthy donor, CLL and MDS specimens were obtained from healthy volunteers or patients cared for at Stanford University under institutional review board (IRB) protocols #18329 and #71851 and analyzed under IRB protocol #65904. Both male and female donors were used. All participants provided written informed consent. *SF3B1* mutations were determined using Sanger sequencing, HemeSTAMP next generation sequencing (NGS)^69^ or the Sequentify™ panel^70,71^. Unless otherwise stated, peripheral blood mononuclear cells (PBMCs) were studied. No randomization was performed.

#### Cell Culture

Human K562 cells (CCL-243) and NALM6 cells (CRL-3273) were obtained from the American Type Culture Collection. BLCL, SC-1, HEK293T, Raji and MEC-1 cells were a kind gift from Dr. Christine Mayr’s laboratory at Memorial Sloan Kettering Cancer Center. K562 cells were maintained in Iscove’s Modification of DMEM (Corning, USA 10-016-CV) supplemented with 10% (*v*/*v*) FBS (Gibco) and 1% penicillin/streptomycin (P/S) (Cytiva, USA, SV30010). BLCL, SC-1, Raji, and NALM6 were cultured in RPMI-1640 medium (Corning, USA, 10-041-CV) containing 10% FBS and 1% P/S. MEC-1 cells were cultured in RPMI-1640 media containing 20% FBS and 1% P/S. HEK293T cells were cultured in DMEM (Sigma, Aldrich, USA, D5796) supplemented with 10% FBS and 1% P/S. HCT116 WT and p53 null cell lines were a kind gift from Dr. Bert Vogelstein’s laboratory at Johns Hopkins University and were cultured in McCoy’s 5A media (Corning, USA, 10-050-CV) supplemented with 10% FBS and 1% P/S. All the cells were incubated in a humidified atmosphere at 37°C and 5% CO_2_. All cells were periodically authenticated by MD Anderson’s Cytogenetics and Cell Authentication Core.

#### Mutant Cell Line Generation

Lentiviruses bearing shRNAs were generated using 293T cells. Briefly, 293T cells were transfected with pLenti-TRC2 plasmids containing shRNAs, as well as viral protein plasmids dR8.2 (Addgene #12263) and pCMV-VSV-G (Addgene #8454). Viruses were harvested 48 to 64 hrs after transfection and concentrated with Retro X (Takara Bio, 631456). Retroviruses were generated by transfecting 293T cells with pMSCV-PIG plasmids containing IPADP1, pCMV-VSV-G and pMVC (a kind gift from Dr. Christine Mayr’s laboratory at MSKCC).

To generate stable IPADP1 knockdown and control cell lines, K562, NALM6, and MEC-1 cells were transduced with lentiviruses carrying shRNAs in the culture plates coated with 0.05 mg/mL poly-D-lysine for 1 hr at 37°C. Cells were spin-infected containing 10 μg/ml polybrene (Sigma TR-1003) at 933 x *g* at 33°C for 45 min. Transduced cells were sorted using fluorescence-activated cell sorting (FACS) with a CytoFLEX SRT sorter (Beckman, USA) 5 days after infection. To generate ectopically expressed IPADP1 or *p53-IPA* cells, K562 and HCT116 cells were transduced with retrovirus containing 10 μg/ml polybrene. After cultivating for another two days, the infected cells were selected using 0.5 μg/ml puromycin. To generate IPADP1-mCherry expressing cells, HEK293T cells were transfected with Lipofectamine 3000 (Invitrogen, USA, L3000015) according to the manufacturer’s protocols.

#### Constructs

For the generation of shRNA-mediated stable IPADP1 knockdown, pLenti-TRC2 vector was first edited to have a KpnI site and then digested with KpnI and EcoRI. shRNA oligonucleotides were obtained from Intergrated DNA Technologies (IDT, USA), annealed, and ligated with vector using T4 ligase. After transformation, clones were screened through DNA extraction and Sanger sequencing. shRNA oligonucleotides can be found in Table S4.

For ectopic expression of IPADP1 and *p53-IPA*, pMSCV-PIG was obtained from Addgene (#21654). Flag tag was cloned into pMSCV-PIG with BglII and XhoI. IPADP1 (ORF1) and ORF2 fragments were amplified from BLCL genomic DNA and cloned with XhoI and EcoRI. Full-length *p53-IPA* was amplified from BLCL genomic DNA and cloned with BglII and XhoI. The fusion protein of mCherry and IPADP1 was generated by Gibson cloning. The mCherry fragment was amplified from pcDNA3.1-mCherry (Addgene #128744) and IPADP1 fragment was amplified from the pMSCV-PIG-IPADP1 plasmid described above. Constructs were transformed and sent out for sequencing as described above. The primers used for amplification of the pieces are listed in Table S4.

All gene cloning, manipulation, and plasmid propagation steps involving pcDNA3.1, pLenti-TRC2, pMSCV-PIG, ZeroBlunt, and viral protein plasmids were carried out in *Escherichia coli* 5-alpha (New England Biolabs), TOP10 (Invitrogen, C404010) or *Escherichia coli* NEB Stable cells (New England Biolabs, USA, C3040H) grown in LB media supplemented with appropriate selection antibiotics.

#### CRISPR

CRISPR was performed in K562 largely following IDT protocol for Alt-R CRISPR-cas9 electroporation delivery of RNP. Custom Alt-R crRNA was designed against IPA sequences and combined with Alt-R tracr RNA to form a sgRNA duplex. 3.125 μL of the sgRNA was combined with 25 μg of cas9-GFP (IDT, USA, 10008161) and electroporated into K562 cells using the Lonza SF nucleofection kit (USA, V4XC-2032) and Lonza 4-D Nucleofector (USA, AAF-1003B) according to the manufacturer’s instructions. Electroporation code used was FF-120. After electroporation, cells were seeded onto 48-well plates and expanded, after which GFP-expressing cells were sorted singly into 96-well plates by fluorescence-activated cell sorting (FACS) with a CytoFLEX SRT sorter (Beckman, USA). Individual single-cell clones were subsequently expanded and genotyped via Sanger sequencing.

#### Cell Viability Assay

Cells were seeded into 96-well white-walled clear-bottom plates at a range of 3,000 to 30,000 cells per well, depending on the drug treatment, in 100 μL of culture media. On the same day, cells were treated with doxorubicin or cisplatin by adding 50 μL of culture media with appropriate drug concentrations to the wells. Cells were treated with imatinib or dasatinib 24 hrs after plating the cells as described above. Cells were treated with 5-FU 48 hrs after plating as described above. Cells with doxorubicin or cisplatin were treated for 24 hrs, cells with imatinib or dasatinib were treated for 48 hrs, and cells with fludarabine or 5-FU were treated for 72 hrs. After treatment time, Cell Titer Glo (Promega, USA, G9243) was added in a 1:1 ratio to total volume in the well. Using the GloMax Discover Microplate Reader (Promega, USA, GM3000), the plate was orbitally shaken for 2 min at 300 rpm and then incubated in the machine at room temperature for 10 min. After this, luminescence was measured and analyzed using Microsoft Excel and Graphpad Prism 10. First, the luminescence results were normalized to wells containing media without cells, then the results were normalized to the average luminescence of the untreated cells. Each experiment was repeated independently at least three times with triplicates. Data was analyzed using GraphPad Prism software.

#### Apoptosis Assay with Annexin V/DAPI Staining and Flow Cytometry

Annexin V APC (BD Pharmingen, USA, 550475) and DAPI were used to detect cellular apoptosis. Cells were seeded in 48-well plates at a density of 1.2 × 10^5^ cells per well. Cells were immediately treated with doxorubicin (MCE, USA, HY-15142) and cisplatin (Millipore Sigma, USA, 232120) for 24 hrs and imatinib (MCE, USA, HY-15463) for 48 hours. After treatment, cells were harvested in a microcentrifuge tube. After one wash with pre-chilled PBS, cells were suspended in 150 μL binding buffer. 1.5 μL of 2.0 μg/mL annexin V APC and 1.5 μL of 10 μg/mL DAPI was added to each tube and stained for 15 min protecting from light before starting flow cytometry. Finally, detection of staining was performed by Beckmann-Coulter Cytoflex LX Flow Cytometry (Beckmann-Coulter, USA, C40323) and data was analyzed using FlowJo 10.

#### Analysis of γH2AX Expression by Flow Cytometry

Control and IPADP1-KD K562 cells were treated with 5 μM doxorubicin (MCE, USA, HY-15142) for 1 hr, then drug was removed via centrifugation at 500 × *g* for 5 min at room temperature. Fresh media was added back, and cells were plated in a 6-well plate with control cells and IPADP1-KD K562 cells at cell densities of 5 × 10^5^ cells per well and 1 × 10^6^ cells per well respectively. At each recovery timepoint, cells were fixed with 4% formaldehyde (Electron Microscopy Sciences, USA, Cat. #15710) for 15 min in the dark at room temperature. Fixed cells were washed once with sterile PBS then frozen down at -80°C with freezing media (90% FBS, 10% media). On the day of flow cytometry analysis, frozen cells were first thawed with 500 μL chilled FACS buffer (sterile PBS containing 2% FBS) without resuspension. Cells were spun down at 1000 × *g* for 5 min at room temperature and washed once with FACS buffer. Cells were permeabilized with pre-chilled 90% methanol and incubated on ice for 5 mins, then washed twice with FACS buffer. Control and IPADP1-KD K562 cells were then re-hydrated with FACS buffer. The volumes of both control and IPADP1-KD cells were equally divided into unstained and stained groups. The stained group was stained with H2AX (BD Pharmingen, USA, Cat. 560447) and KI67 (BD Pharmingen, USA, Cat. 561283). Cells in the stained group were stained in the dark for 30 min on ice, washed once with FACS buffer, then resuspended in 200 μL of FACS buffer. Detection of staining was performed in a Costar 96-well round bottom plate (Corning Inc, REF 3799) by Beckmann-Coulter Cytoflex LX flow cytometry and data was analyzed using FlowJo 10 software.

#### Cell Cycle Assay by EdU and DAPI Labelling and Detection with Flow Cytometry

K562 and HCT116 cells were seeded in 48-well plates at a density of 2 × 10^5^ cells per well. Cells were immediately treated with doxorubicin (MCE, USA, HY-15142) for 0, 24, and 48 hrs. After 24 hrs treatment, cells were spun down at 500 x *g* for 5 minutes and resuspended in fresh media. For EdU labelling using AF647 click-EdU kit (BD Pharmagen, USA, 565456), 5 μM EdU or DMSO was added to basal medium of K562 for 2 hrs prior to harvesting. Cells were pelleted, washed once with PBS, and fixed in 4% paraformaldehyde (PFA) in PBS for 15 min at room temperature. PFA was washed off cells with two rinses with PBS and resuspended in fetal bovine serum (FBS) (Gibco, USA, A56708-01) + 12% DMSO. Cells were then frozen at -80°C for at least 15 min. On the day of EdU labeling and flow cytometry, cells were permeabilized with a 10 min incubation in kit-provided saponin permeabilization buffer at room temperature, followed by a 30 min incubation in click-EdU reaction buffer using azide-AlexaFluor-647. Cells were rinsed twice with PBS, then resuspended in 2% FBS in PBS. 6.5 μg/mL DAPI was added and incubated for 15 minutes at room temperature. Staining was detected by Beckmann-Coulter Cytoflex LX flow cytometry and data was analyzed using FlowJo 10.

#### Western Blotting

Cells were lysed on ice for 10 min with lysis buffer (30 mM Tris pH 7.4, 150 mM NaCl, 0.5% Triton X-100, 1 mM EDTA, 0.05% SDS), containing freshly added protease inhibitor (Thermo Scientific, USA, 1861278) and phosphatase inhibitor cocktails (Thermo Scientific, USA, P5726 and P0044). Lysates were boiled with 6x Laemmli buffer at 75°C for 10 min. Any lysates intended to be blotted for IPADP1 were boiled at 70°C for 12 min. Lysates were run on 4–12% bis-tris NuPAGE gels (Invitrogen, USA, NP0336BOX) with MES running buffer (Invitrogen, USA, NP0002). The separated proteins were transferred to nitrocellulose membranes (Bio-Rad, USA, 1620252), blocked with Intercept PBS blocking buffer (Li-Cor, USA, 927-70001) for 1 hr at room temperature, followed by incubation with primary antibodies at 4°C overnight. Any Western blot for IPADP1 was blocked with primary antibodies at 4°C for 2-3 days. After three washes using PBS with 0.1% tween-20 (PBST), the blots were incubated with IRDye-conjugated secondary antibodies for 1 h at room temperature. Any western blot for IPADP1 had 4 washes with PBST. After three washes with PBS, proteins were detected with a LiCor Odyssey F imaging system. Any western blot for IPADP1 had 4 washes with PBS before proteins were detected using the Odyssey F imaging system.

The following primary antibodies were used: anti-Flag (mouse, Sigma, F3165), anti-Beta Actin (rabbit, ProteinTech, 20536-1-AP; mouse, Sigma, A2228), anti-Histone H3 (rabbit, Cell Signaling, 14269T), anti-BAG6 (mouse, Santa Cruz Biotechnology, sc-365928), anti-p53 (mouse, Santa Cruz Biotechnology, sc-126), anti-p21 (rabbit, ProteinTech, 10355-1-AP), anti-MDM2 (rabbit, Cell Signaling Technology, 86934S), anti-SRP54 (mouse, Santa Cruz Biotechnology, sc-393855), anti-Bax (rabbit, Cell Signaling Technology, 5023S), anti-PARP (rabbit, Cell Signaling Technology, 9542S), anti-Pro-Caspase 3 (mouse, Cell Signaling Technology, 9668T), anti-Cleaved Caspase 3 (rabbit, Cell Signaling Technology, 9661T), anti-UMPS (mouse, Santa Cruz Biotechnology, sc-398086), anti-CDK1 (rabbit, ProteinTech, 10762-1-AP), anti-AIFM1 (rabbit, ProteinTech, 17984-1-AP), anti-DNM1L (rabbit, ProteinTech,26187-1-AP), anti-MSH2 (rabbit, ProteinTech,15520-1-AP). The secondary antibodies used included anti-rabbit IRDye 680 (donkey, Li-Cor Biosciences, 926-68073), anti-rabbit IRDye 800 (donkey, Li-Cor Biosciences, 926-32213), anti-mouse IRDye 800 (donkey, Li-Cor Biosciences, 926-32212), and anti-mouse IRDye 680 (donkey, Li-Cor Biosciences, 926-68072).

The custom rabbit polyclonal antibody for IPADP1 was produced by GenScript using a combined injection of 3 peptides, followed by purification of the antibody using all 3 peptides. The peptide sequences are: CGSRLHFGRPRRAD, GVPDQRGQRGESP, IREGEAGESLEPGR.

#### Western Blot with Drug Treatments

HCT116 cells were seeded in 12-well plates at a density of 1.25 x 10^5^ cells per well and grown for 32 hrs before addition of concentrations of 5-fluorouracil (5-FU) ranging from 0 to 200 µM. Cells were harvested 6, 16, and 24 hrs later and then western blot analysis was performed as described above. HCT116 cells were seeded in 12-well plates at a density of 1.25 x10^5^ cells per well. 16 hrs after plating 5 µM 5-FU was added. Cells were harvested at hours ranging from 0 to 72 hrs after treatment and western blot analysis was performed as described above.

K562 cells were seeded in 12-well plates at a density of 5 x 10^5^ cells per well and immediately 2.5 or 5 µM doxorubicin or 5 µM cisplatin was added. Cells were harvested 16 hrs later and western blot analysis was performed as describe above.

#### Cell Fractionation

K562 cells were washed with cold PBS once and lysed with lysis buffer (10 mM Tris/Cl pH 7.5, 150 mM NaCl, 0.5 mM EDTA) containing 0.019% NP-40 for 5 seconds on ice. Lysates were spun down at 21,000 x *g* for 8 mins at 4°C. Supernatant was used as the cytoplasm fraction, and pellet was the nuclear fraction. Lysates of each fraction were mixed with 6x Laemmli sample buffer and performed western blot as described above.

### Protein Crosslinking Western Blot

HCT116 cells with vector control and IPADP1 expression vector were seeded in 6-cm culture dishes at a density of 2 million cells per dish and grown for 32 hrs before treating with 10 μM 5-FU for 16 hrs. Cells were harvested and lysed in 200 μL of crosslinking lysis buffer (50 mM Tris-HCl pH=8, 150 mM NaCl, 1 mM EDTA pH=8, 1% (v/v) NP-40) with fresh protease and phosphatase inhibitors. Cells were placed on a shaker at 4°C for 30 min at 500 rpm and then centrifuged at 13,000 rpm for 10 min at 4°C. Lysates were aliquoted into volumes of 25 μL and glutaraldehyde ranging from 0.01% to 0.05% (v/v) were added to the samples and shaken for 5 min. After that, samples were inactivated using 6x Laemmli sample buffer with SDS (Boston Labs, USA, BP-111R). Crosslinked samples were used for western blotting following the protocol described above.

#### Immunoprecipitation

MSCV-PIG Ctrl and MSCV-PIG Flag-IPADP1 were transfected into HEK293T cells using Lipofectamine 3000 (Invitrogen, USA, L3000015). After 24 hrs, transfected cells were collected and lysed on ice for 30 min with Fab trap lysis buffer (Chromotek) containing freshly added protease inhibitor cocktail (Thermo Scientific) and phosphatase inhibitors (Sigma). After centrifugation at 21,130 × *g* for 10 min, Flag-Fab Trap immunoprecipitation was performed following the manufacturer′s instructions. In brief, Flag-Fab Trap slurry (Chromotek) was added, incubated with cell lysate for 2 hrs at 4°C on a rotator, and washed three times with pre-chilled dilution buffer. Washed slurry was mixed with 6x Laemmli sample buffer and boiled at 95°C for 5 min or 75°C for 10 min. The immunoprecipitates were run on 4-12% Bis-Tris NuPAGE gels using MES running buffer, followed by western blot analysis of the endogenously expressed candidate interactors.

MSCV-PIG Ctrl and MSCV-PIG Flag-IPADP1 expressing HCT116 cells were collected and lysed on ice for 30 min with lysis buffer (Chromotek) containing freshly added protease inhibitor cocktail (Thermo Scientific) and phosphatase inhibitors (Sigma). After centrifugation at 21,130 × *g* for 10 min, p53 N-term Trap agarose (Chromotek, pta-20) and p53 C-term Trap agarose (Chromotek, pta2-20) immunoprecipitation was performed following the manufacturer′s instructions. Western blot analysis was performed as described above.

#### Sample Preparation for Mass Spectrometry Analysis

After fab Trap co-IP described as above, the pull-down protein solution was run on a gel following by SimplyBlue staining (Invitrogen, USA, 465034) and submit to Taplin Mass Spectrometry Facility at Harvard Medical School for mass spectrometry analysis.

#### Gene Ontology Analysis

Downstream Gene Set Enrichment Analysis was performed utilizing Fisher Exact test. Proteins that are significantly differential (*P<0.05*) in the treatment compared to the control group were tested based on MSigDB (https://www.gsea-msigdb.org/gsea/msigdb/collections.jsp). Hallmark and C2 curated gene sets. *P* values are adjusted for multiple testing using Bonferroni method.

#### Clonogenic Assay

HCT116 cells with vector control and IPADP1 expression were seeded in 6-well plates at a density of 1000 cells per well in culture media. After 24 hrs, cells were treated with 1 and 5 μM 5-FU for 48 hrs. Cells were grown for approximately 3 weeks until colonies were visible, with culture media changed every 3 days. At collection point, colonies were stained with methylene blue for 30 min at room temperature, washed three times with PBS, and then dissolved with 1% N-lauroyl sarcosine in water. The plates were incubated in 1% N-lauroyl sarcosine for 30 min on a shaker and then moved to a 96-well microplate in triplicates. Absorbance at 600 nm was read using the Promega Glomax Discover.

#### Fluorescent Microscope Imaging

Fluorescent images were taken with a Zeiss Axio Observer 5 (USA, 491916-0001-000) containing 405 nm, 488 nm, and 561 nm lasers. Images were captured and analyzed with Zeiss Zen Version 3.1.1 Software.

#### Protein Structure Prediction

All protein structure prediction was done using ColabFold v1.5.5: AlphaFold2 using MMseqs2.

#### Xenograft Study

All animal studies were conducted in compliance with institutional guidelines and were approved by the Animal Welfare Committee of the University of Texas Health Science Center at Houston (protocol #AWC-24-0108). Female nude mice (7–8 weeks old) were purchased from The Jackson Laboratory and acclimated before experimentation. To examine the role of IPADP1 in leukemia tumorigenesis, Matrigel (Corning, USA, 356234) was diluted 1:1 with Iscove’s Modification of DMEM medium to prepare a 50% solution. A total of 1 × 10^6^ cells (K562 shCtrl, shIPADP1-1, shIPADP1-2) were then mixed with the 50% Matrigel at a 1:1 ratio and injected subcutaneously into the right hind flank of each mouse. Tumor growth was assessed twice weekly by caliper measurements of tumor length and width. After 40 days, mice were euthanized, and tumors were harvested for further analysis. Tumor volume was calculated using the formula: (length × width²) / 2.

#### RT-PCR and qRT-PCR

Total RNA was isolated using Monarch RNA extraction Kit with DNase treatment (New England Biolabs T2110S) following manufacturer’s instructions. RNA was reverse transcribed using the qScript cDNA SuperMix (QuantaBio 95048-100) or the SuperScript IV First-Strand Synthesis System (Invitrogen 00745157). RT-PCR reactions were carried out using Q5 high-fidelity DNA polymerase (New England Biolabs, M0491) with manufacturer’s protocol. qRT-PCR reactions were carried out using Powertrack Sybr Green Master Mix (Applied Biosystems, USA, C14512) on a CFX384 Touch Real-Time PCR Detection System (Bio-Rad, USA, 1855484). Primers were designed to be intron-spanning and are listed in Table S4. Primers that recognized genomic DNA and pre-RNA products were included to control *p53-IPA* expression.

#### Immunofluorescence

Cells were plated on 4-well or 8-well Millicell EZ slides (Millipore, PEZGS0496, PEZGS0816). The next day, cells were washed with PBS, fixed in 4% paraformaldehyde for 10 min at room temperature (RT) and washed 2 x 5 min with PBST (PBS containing 0.1% Tween). Then, cells were permeabilized in 0.1% Triton X-100 in PBS for 15 min at RT and blocked with blocking buffer (3% BSA in PBS + 0.1% Tween) for one hour and gently rocked at RT. After that, cells were incubated in the diluted primary antibody in blocking buffer for 2 hours at RT. Cells were then washed 3 x 5 min with PBST and incubated with the secondary antibody in blocking buffer for 1 hour at RT in the dark. After three washes using PBST, the slide was mounted with ProLong^TM^ Gold Antifade Mountant with DNA Stain DAPI (Invitrogen, P36931) and a coverslip was added. Images were captured using a Zeiss Axio Observer 5 and analyzed with Zeiss Zen Version 3.1.1 Software.

#### RNA extraction and QuantSeq-REV 3′ mRNA sequencing

RNA was extracted using Trizol™ (cat. #15596026) from ThermoFisher Scientific (Waltham, Massachusetts) as per manufacturer recommendations and quantified via Nanodrop and/or Qubit using the RNA broad range assay kit (cat. #Q10211). RNA was further analyzed for quality using an Agilent Bioanalyzer with the RNA 6000 Nano kit (cat. #5067-1511). Libraries were prepared using the QuantSeq REV kit (cat. 16.96) with PCR add-on kit (cat. 20.96) from Lexogen GmbH (Vienna, Austria) per manufacturer’s instructions. Samples were barcoded using the Lexogen i5 6 nt dual indexing add-on kit (Cat. 47.96), and underwent quantification, pooling, and sequencing on Illumina HiSeq X Ten or Novaseq X instruments to generate pair-end 150 bp reads by Novogene Inc. (Sacramento, CA). As QuantSeq REV requires a custom sequencing primer, no PhiX spike-in was used. Sequencing depth generally exceeded > 20 million reads per specimen.

#### Quantification of *p53-IPA* expression and its association with *SF3B1* mutation status

QuantSeq-REV 3’ mRNA sequencing reads were processed and aligned using the APAlyzer pipeline (https://github.com/RJWANGbioinfo/APAlyzer/). Briefly, raw reads were trimmed with BBDuk (BBTools v39.01) to remove 3’ adapter sequences and low-quality bases (minimum length 20 bp), then aligned to the hg38 reference genome using STAR (v2.7.11b) with standard splicing-aware parameters. Aligned reads were intersected with annotated polyadenylation cleavage sites from the PolyA_DB v3 database (http://polya-db.org/polya_db/v3) to quantify APA isoform usage across all genomic compartments, encompassing 3’ UTR, coding sequence (CDS), and intronic APA (IPA) sites. To account for positional uncertainty in cleavage site annotation and read mapping, each reference site was extended by a symmetric flanking window of ±24 bp and reads overlapping this interval were assigned as supporting evidence for the corresponding polyadenylation events. Per-sample IPA usage ratios were calculated as the fraction of reads supporting specific IPA site (chr17:7685249) relative to total reads mapping to *TP53* gene (chr17:7661779-7687550).

##### CLL analysis

Normal B cells (n=5), which represent the non-malignant cellular compartment of origin for CLL (n=29), were used as the reference group. The mean and standard deviation of IPA usage were computed across B-cell samples, and a significance threshold was set at mean + 2SD. A diluted Beta prior was constructed from the reference group by pooling site-supporting reads and total *TP53*-mapping reads across all reference samples, scaled by a dilution factor of 0.05 to allow data-driven updating while anchoring the prior to the B-cell baseline. A Bayesian model with a Beta-Binomial likelihood and conjugate Beta prior was applied to classify samples as *p53-IPA* positive. For each test sample, the posterior probability that true IPA usage exceeded the reference-derived threshold was computed using the Beta cumulative distribution function: P(θ > threshold | data) = 1 − pbeta(threshold, α₀ + k, β₀ + n − k) where k is the number of IPA site-supporting reads, n is the sum of reads mapping to *TP53*, and α₀ and β₀ are the prior shape parameters. Samples with a posterior probability >95% were classified as highly positive, and samples with posterior probability >50% but ≤95% were classified as lowly positive. Samples were required to have a minimum of 10 total *TP53*-mapping reads and at least 1 site-supporting read to be eligible for classification. All reference samples returned a posterior probability below the positive threshold, confirming zero false-positive calls in the normal reference groups.

##### MDS analysis (n=65)

*SF3B1*-wildtype (SF3B1-WT) MDS samples (n=31) were used as the reference group. Prior to threshold setting, the WT reference distribution was inspected for outliers using the Tukey IQR method (upper fence = Q3 + 1.5×IQR), which identified two WT samples (S681, S686) with disproportionately elevated IPA usage and were excluded from the reference to avoid inflating the baseline. The significance threshold was set at the clean WT mean + 1SD, which yields zero false-positive calls among clean WT samples. A diluted Beta prior (reduced dilution factor of 0.01 applied to allow individual sample evidence to dominate over the prior) was constructed by pooling site-supporting reads and total *TP53*-mapping reads across the remaining clean WT reference samples. Same parameters as above were applied to classify samples with *p53-IPA* expression. Samples with a posterior probability >95% were classified as positive.

The association between *p53-IPA* positivity and *SF3B1* mutation status was evaluated in the CLL and MDS cohorts, respectively. Samples were dichotomized as IPA-positive or IPA-negative and compared against *SF3B1* mutation status in a 2×2 contingency table. Odds ratios (OR) were estimated thereof, representing times more likely *p53-IPA* positivity is to occur in *SF3B1*-mutant samples compared to *SF3B1*-wildtype samples. Statistical significance was assessed using Boschloo’s exact test conditioned on the fixed *p53-IPA* positive margin. All analyses were performed in R (4.3.1).

#### Statistics

All statistics were performed using GraphPad Prism 10.3.1 software. Unless otherwise specified, unlabeled denotes no significance, * denotes *P* < 0.05, ** denotes *P* < 0.01, *** denotes *P* < 0.001, and **** denotes *P* < 0.0001. Statistical analyses were calculated using Welch’s t-test.

---

**Figure S1.**
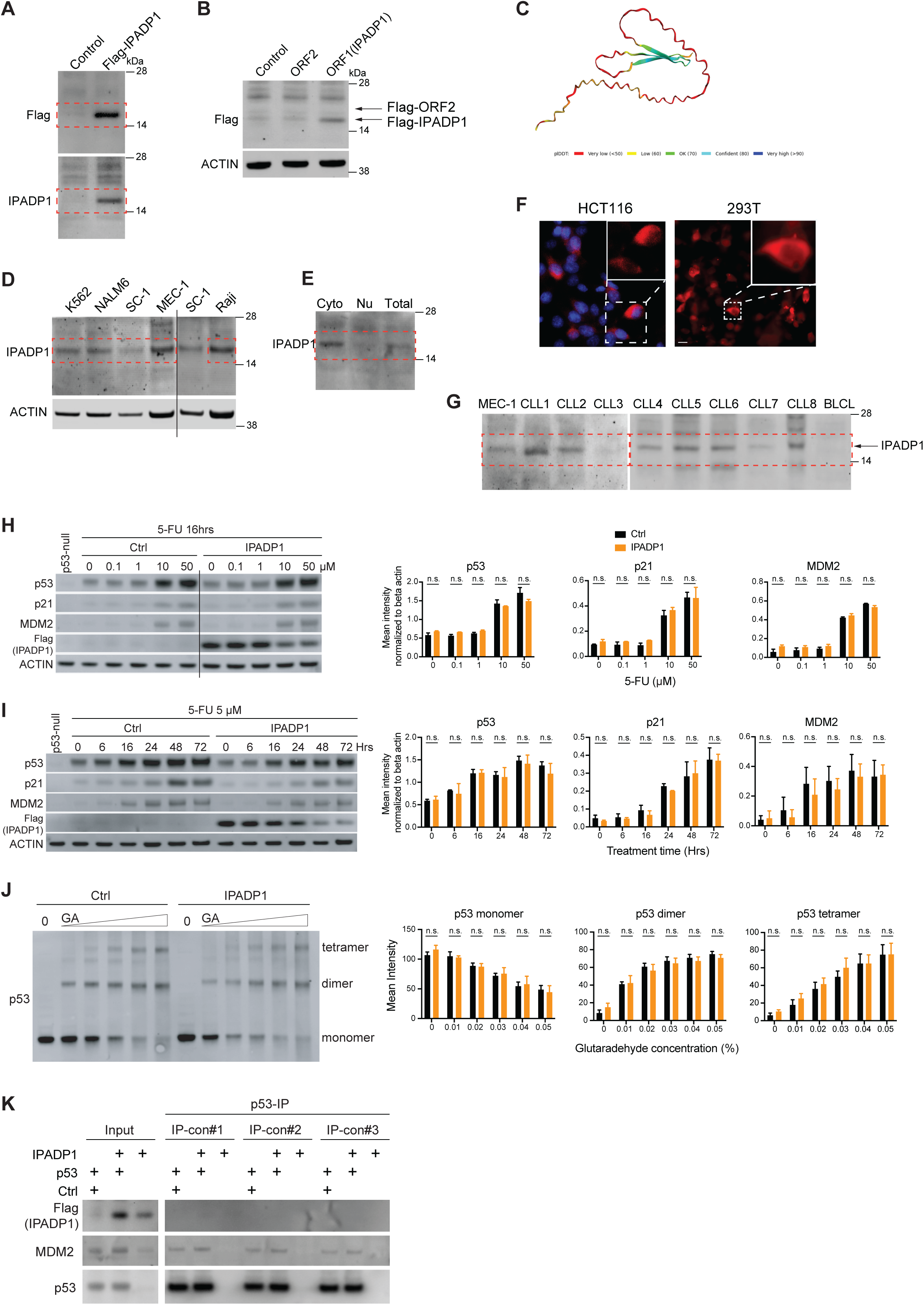
***p53-IPA* generates a non-canonical protein, IPADP1 (A)** The uncropped image of Western blot result in Figure 1B. **(B)** Western blot analysis in HCT116 cells expressing two putative Flag-tagged open reading frames (ORF). ORF1 is IPADP1. ACTIN, loading control. **(C)** ColabFold prediction of protein structure of IPADP1. **(D)** The uncropped image of Western blot result in Figure 1D. **(E)** The uncropped image of Western blot result in Figure 1E. **(F)** Left, representative fluorescence image of HCT116 cells transfected with Flag-tagged IPADP1 and stained with an anti-Flag antibody. Blue: DAPI. Red: Flag-IPADP1. Right, representative fluorescence image of HEK293T cells transfected with IPADP1 fused with mCherry. Scale bars = 20 µm. **(G)** The uncropped image of IPADP1 expression for Figure 1 F. IPADP1 band is indicated. The same band was shown decreased in MEC-1 cells with shRNA knockdown (Fig S2A). **(H)** Western blot analysis in HCT116 cells, with the expression of control vector or IPADP1, treated with varying concentrations of 5-fluorouracil (5-FU) for 16 hrs. p53-null HCT116 cells was used as a negative control for p53, p21, and MDM2 expression. ACTIN, loading control. No significant difference in p53 accumulation between control (Ctrl) and IPADP1 expression was observed. The quantification is shown on the right (N = 2). n. s., non-significant. **(I)** As in (E), but cells were treated with 5 µM of 5-FU for varying timepoints from 0 to 72 hrs. No significant difference in p53 accumulation between control (Ctrl) and IPADP1 expression was observed. The quantification is shown on the right (N = 2). n. s., non-significant. **(J)** Determination of p53 oligomerization status using glutaraldehyde (GA) cross-linking. HCT116 cells with control plasmid or IPADP1 expression were treated with 10 µM 5-FU for 16 hrs and lysed. Protein lysates were treated with increasing concentrations of GA for 5 mins, after which western blot analysis was performed with an anti-p53 antibody. Bands show monomers, dimers, and tetramers of p53. Quantified results, N = 5. No significant difference in p53 oligomerization between control (Ctrl) and IPADP1 expression was observed. n. s., non-significant. **(K)** Assessed the association of IPADP1 and p53 protein using protein immunoprecipitation (IP) coupled with Western blot analysis in HCT116 cells. Cells were treated with 5 µM of 5-FU for 16 hrs to induce endogenous p53 protein accumulation. p53 protein was pulled down with p53 agarose beads with three IP conditions-#1, the beads were washed with less stringent wash buffer 3 times; #2, same wash buffer as #1 but wash 4 times; #3, the beads were washed with wash buffer containing 0.5% NP-40 3 times. This is shown as a representative result of 2 repeats.

**Figure S2.**
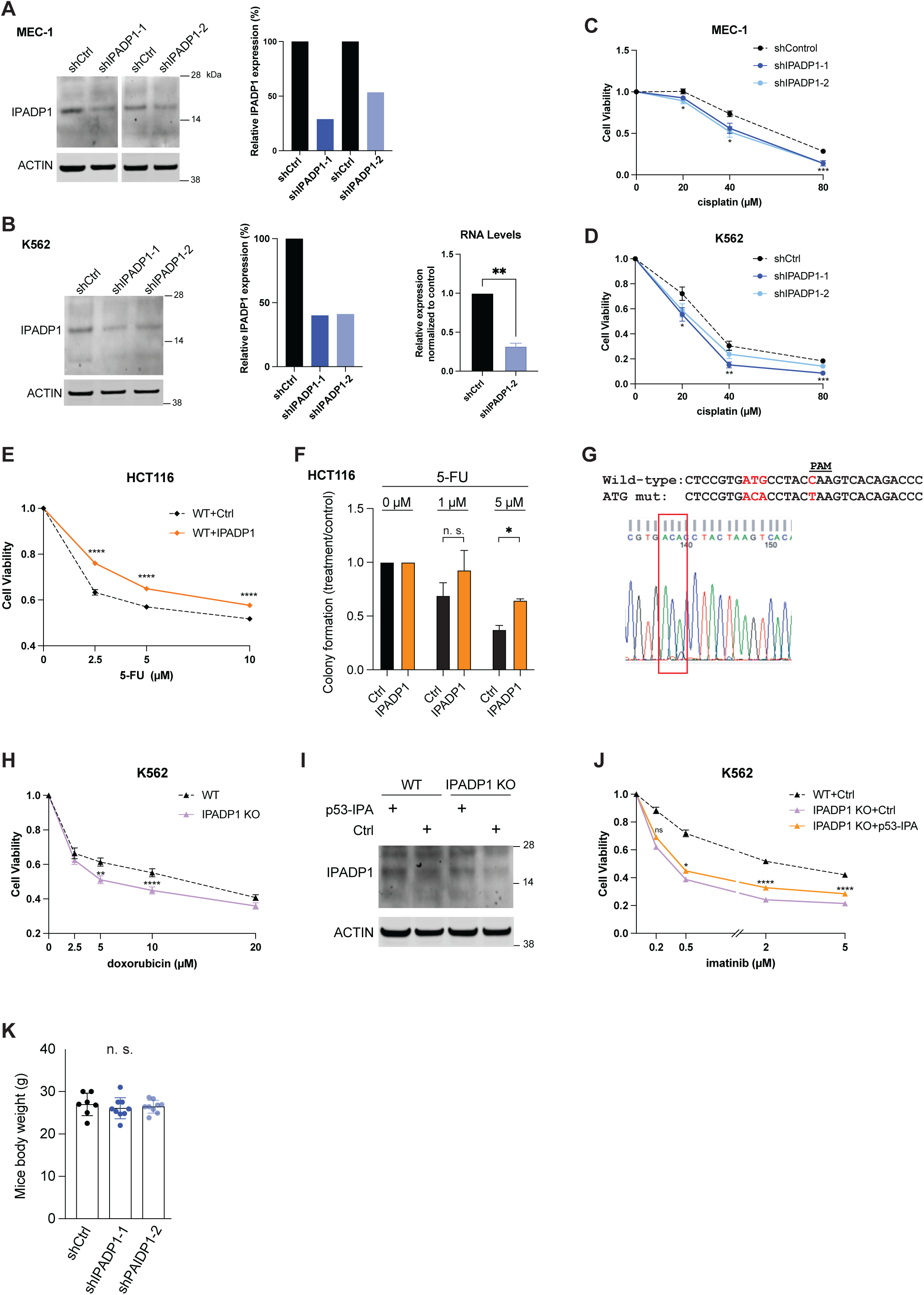
IPADP1 affects cell viability upon drug treatments. **(A)** Western blot analysis of endogenous IPADP1 in MEC-1 shRNA-mediated IPADP1-knockdown and control cells. ACTIN, loading control. The quantification is shown on the right. **(B)** Western blot analysis of endogenous IPADP1 in K562 shRNA-mediated IPADP1-knockdown and control cells. ACTIN, loading control. The quantification is shown on the right. The knockdown of shIPADP1-2 was further confirmed by qPCR. N = 3. **, *P* < 0.01. **(C)** As in Fig 2A but performed with MEC-1 control and IPADP1 knockdown cells treated with cisplatin for 24 hrs. N = 3 biologically independent experiments. **(D)** As in Fig 2A but performed with K562 control and IPADP1 knockdown cells treated with cisplatin for 24 hrs. N = 3 biologically independent experiments. **(E)** As in Fig 2A but performed with HCT116 cells with vector control or IPADP1 ectopic expression treated with 5-FU for 72 hrs. N = 3 biologically independent experiments. **(F)** The clonogenic assay of HCT116 cells with vector control or IPADP1 ectopic expression treated with 5-FU. Cells were plated at a cell density of 1000 cells per well and treated with 5-FU for 48 hrs. Data was first normalized to a blank and then to untreated values within cell lines. \**P* < 0.05. n. s., non-significant. N = 3 biologically independent experiments. **(G)** Sanger Sequencing results (top) and chromatogram (bottom) for K562 cells CRISPR edited to mutate translation start codon (labeled in red) of IPADP1. **(H)** As in Fig 2A but performed with K562 wild-type (WT), IPADP1 CRISPR knockout (IPADP1 KO) cells treated with doxorubicin for 24hrs. N = 3 biologically independent experiments. **(I)** Western blot analysis of IPADP1 expression rescue in wild-type (WT) and IPADP1 KO cells. ACTIN, loading control. **(J)** As in Fig 2A but performed with K562 wild-type plus vector control (WT+Ctrl), IPADP1-knockout plus vector control (IPADP1 KO+Ctrl), and IPADP1 rescue cells (IPADP1 KO+p53-IPA) that were treated with imatinib for 48 hrs. N = 3 biologically independent experiments. **(K)** The measurement of mice body weight before tumor excision. n. s., non-significant.

**Figure S3.**
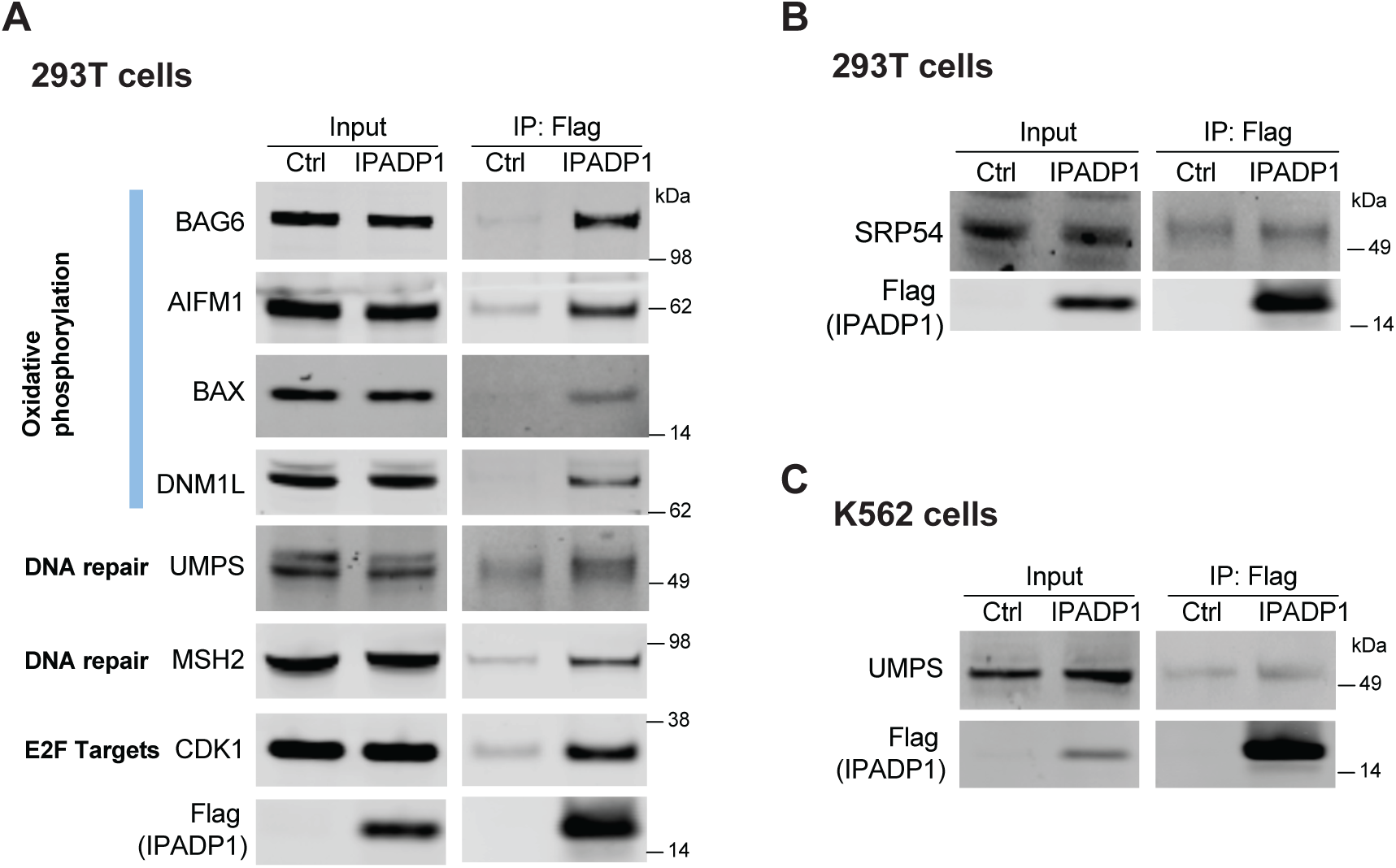
The identification and validation of interactors of IPADP1. **(A)** Western blot validation of IPADP1 interactors in 293T cells. The interactors were identified by Flag protein immunoprecipitation in the control or IPADP1-expressed cells. 0.5% input was loaded. **(B)** As in S3A but analyzed by the SRP54 antibody. **(C)** As in 3B but analyzed by the UMPS antibody.

**Figure S4.**
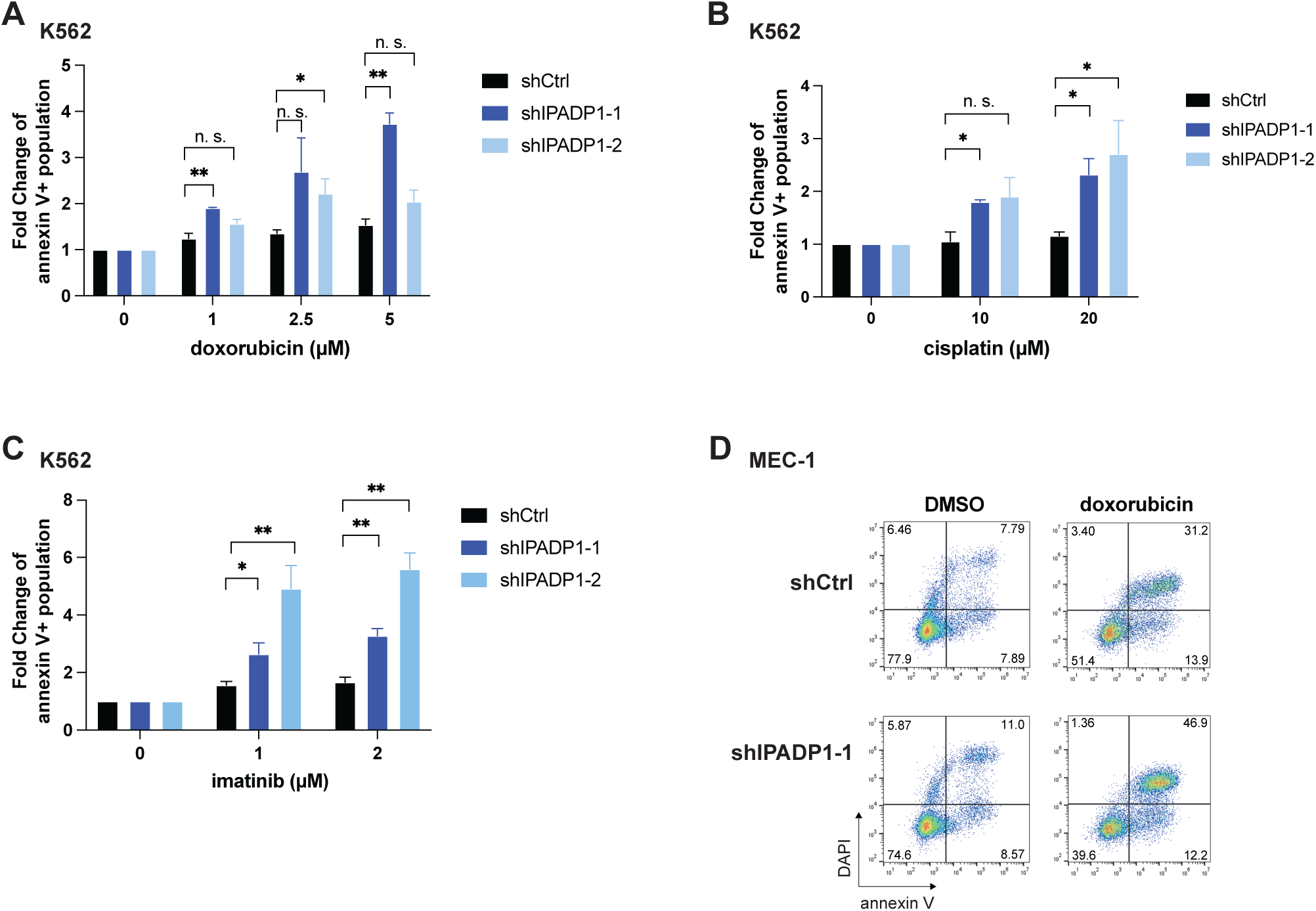
IPADP1 inhibits apoptosis upon drug treatments. **(A)** Fold changes in percentage of annexin V single-positive cells from results of Fig 4A \**P* < 0.05, \*\**P* < 0.01, n. s., not significant. N = 3 biologically independent experiments. **(B)** Fold changes in percentage of annexin V single-positive cells from results of Fig 4B. \**P* < 0.05, n. s., not significant. N = 3 biologically independent experiments. **(C)** Fold changes in percentage of annexin V single-positive cells from results of Fig 4C. \**P* < 0.05, \*\**P* < 0.01. N = 3 biologically independent experiments. **(D)** As in Fig 4A, but performed in MEC-1 cells treated with 0.5 µM doxorubicin for 48 hrs. This is shown as a representative example.

**Figure S5.**
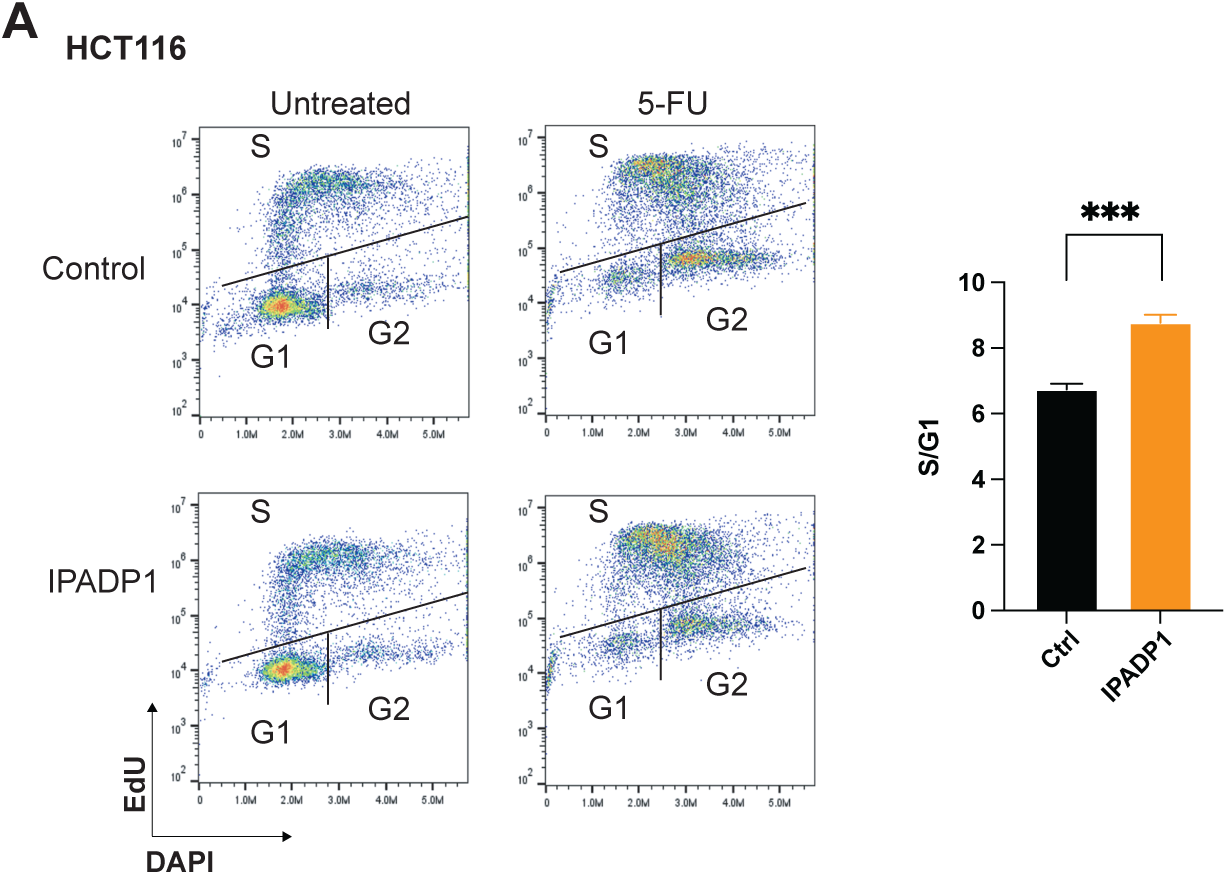
IPADP1 regulates cell cycle. **(A)** As in 5A but done with vector control and IPADP1-expressed p53-null HCT116 cells treated with 5-FU for 72 hrs. \*\*\**P* < 0.001. N = 3 biologically independent experiments.

